# Versatile and automated workflow for the analysis of oligodendroglial calcium signals in preclinical mouse models of myelin repair

**DOI:** 10.1101/2022.10.14.512256

**Authors:** Dorien A. Maas, Blandine Manot-Saillet, Philippe Bun, Chloé Habermacher, Corinne Poilbout, Filippo Rusconi, Maria Cecilia Angulo

**Author notes:** **Corresponding author:** María Cecilia Angulo, Institute of Psychiatry and Neuroscience of Paris (IPNP), INSERM 1266, 102, rue de la Santé, 75014 Paris, FRANCE. These authors contributed equally to this work (Dorien A. Maas, Blandine Manot-Saillet).

## Abstract

Intracellular Ca^2+^ signals of oligodendroglia, the myelin-forming cells of the central nervous system, regulate vital cellular processes including myelination. However, studies on oligodendroglia Ca^2+^ signal dynamics are still scarce, especially during myelin repair, and there are no software solutions to properly analyze the unique Ca^2+^ signal characteristics in these cells. Here, we provide a comprehensive experimental and analytical workflow to acquire and analyze Ca^2+^ imaging data of oligodendroglia at the population and single-cell levels in preclinical mouse models of myelin repair. We report diverse *ex vivo* and *in vivo* experimental protocols to obtain reproducible Ca^2+^ imaging data from oligodendroglia in demyelinated lesions. Importantly, we provide an analytical pipeline containing two free, open source and cross-platform software programs, Occam and post-prOccam, that enable the fully automated analysis of one- and two-photon Ca^2+^ imaging datasets from oligodendroglia obtained by either *ex vivo* or *in vivo* Ca^2+^ imaging techniques. This versatile and accessible experimental and analytical framework, which revealed significant but uncorrelated spontaneous Ca^2+^ activity in oligodendroglia inside demyelinated lesions, should facilitate the elucidation of Ca^2+^-mediated mechanisms underlying remyelination and therefore help to accelerate the development of therapeutic strategies for the many myelin-related disorders, such as multiple sclerosis.

## Introduction

In demyelinating diseases such as multiple sclerosis (MS), myelin sheaths produced by oligodendrocytes (OL) are destroyed by the immune system, resulting in OL death and axonal degeneration and, ultimately, in both physical and neurological disabilities (Duncan & Radcliff, 2016). Upon demyelination, new oligodendrocyte precursor cells (OPCs) and OLs can be generated and partial myelin repair can take place (Franklin & Ffrench-Constant, 2017). However, since remyelination is often incomplete, research tools to investigate oligodendroglial dynamics and roles during the processes of demyelination and remyelination are of great importance for the development of therapeutic strategies.

It is now established that oligodendroglial Ca^2+^ signals translate environmental information into cellular processes such as proliferation, differentiation and myelination but little is known about the origin, dynamics or function of Ca^2+^ signals during myelin repair (Paez & Lyons, 2020; Pitman & Young, 2016; Maas et al., 2021). Recent *in vivo* studies in the zebrafish revealed that both OPCs and OLs are capable of Ca^2+^ signaling (Baraban et al., 2018; Krasnow et al., 2018; Marisca et al., 2020; Li et al., 2022). In mouse brain slices, spontaneous Ca^2+^ activity in OPCs (Balia et al., 2017) and OLs (Battefeld et al., 2019) was shown to be high during postnatal development when the myelination process is still ongoing, and to decrease in OLs as the brain matures (Battefeld et al., 2019). Then, Ca^2+^ signals of OLs are reactivated upon demyelination in the adult mouse brain, suggesting a crucial role for oligodendroglial Ca^2+^ signals in demyelinated lesions (Battefeld et al., 2019). To date, however, only one report has explored OL Ca^2+^ signaling during demyelination in *ex vivo* mouse brain slices (Battefeld et al., 2019) and oligodendroglial Ca^2+^ imaging studies in demyelinated mouse models *in vivo* are completely lacking. Moreover, in the above-mentioned studies, not only were the regions of interest (ROIs) manually chosen, potentially introducing bias in the ROI selection outcome, but also complex Ca^2+^ events with multiple peaks were overlooked even though these events are a hallmark of oligodendroglia (see below).

Several pipelines for the analysis of Ca^2+^ imaging data from neurons and astrocytes in brain slices and *in vivo* have been published. Unfortunately, problems arise when using neuron-oriented programs such as CaImAn and EZCalcium for the analysis of oligodendroglial Ca^2+^ imaging data (Cantu et al., 2020; Giovannucci et al., 2019). For instance, classifiers are used to recognize neurons as round somata of a certain size (Giovannucci et al., 2019), while oligodendroglia possess less well-defined shapes and sizes (Xu et al., 2021). Another important drawback of the available pipelines for neuronal Ca^2+^ imaging is the difference in neuronal and oligodendroglial Ca^2+^ signal dynamics. Indeed, neuronal Ca^2+^ signals last around 100 ms, are characterized by fast kinetics and occur with a high frequency (Chua & Morrison, 2016). This is in stark contrast with oligodendroglia that exhibit complex Ca^2+^ dynamics with signals that last anywhere from several seconds to multiple minutes and that are characterized by slow rise and decay times (our data and Battefeld et al., 2019). Another set of problems may arise when using available Ca^2+^ imaging analysis for astrocytes in oligodendroglia: astrocytes are known to exhibit extensive signal propagation within and between cells and Ca^2+^ signaling pipelines such as AQuA are therefore designed to trace Ca^2+^ events in time and space instead of identifying ROIs (Wang et al., 2019). Because we cannot assume that intra- and inter-cellular signal propagation occurs extensively in oligodendroglia, these pipelines are not immediately suitable for the analysis of oligodendroglial Ca^2+^ signaling. Older ROI-based astrocyte Ca^2+^ imaging analysis packages such as GECIquant and CaSCaDe rely on manual input which may introduce bias, making them not fully desirable (Agarwal et al., 2017; Venugopal et al., 2019).

Here, we provide a comprehensive experimental and analytical workflow to investigate oligodendroglial Ca^2+^ imaging data obtained either *ex vivo* or *in vivo* in two different preclinical mouse models of myelin repair, the lysolecithin (LPC) and cuprizone (CPZ) models. We established reproducible experimental protocols aiming to obtain Ca^2+^ imaging data of oligodendroglia both in brain slices and in freely moving mice expressing genetically encoded fluorescent Ca^2+^ indicators. Next, we developed two free, open source and cross-platform software programs, Occam and post-prOccam (Occam: Oligodendroglial cells calcium activity monitoring), for the fully automated analysis of one- and two-photon Ca^2+^ imaging data from oligodendroglia in demyelinated lesions (GNU GPLv3+ license; code repository at: *https://gitlab.com/d5674/occam*). These highly versatile and accessible tools are suitable for the analysis of Ca^2+^ imaging datasets obtained in diverse preparations, matching all the specific requirements for the monitoring of the unique Ca^2+^ event characteristics of oligodendroglia. Our analytical pipeline thus accelerates the elucidation of Ca^2+^-mediated mechanisms underlying myelin repair and will contribute to the development of therapeutic strategies in myelin-related disorders such as MS.

## Results

### Experimental paradigms to obtain *ex vivo* and *in vivo* oligodendroglial Ca^2+^ signals in demyelinated lesions

The goal of this study was to develop both experimental paradigms and a fully automated analytical workflow for the investigation of one- and two-photon intracellular Ca^2+^ imaging data from oligodendroglia in demyelinated lesions of *ex vivo* and *in vivo* preparations (Fig. 1). We validated our experimental and analytical workflow on data obtained in adult *Pdgfrα*^*CreERT(+/-)*^*;Gcamp6f*^*Lox/Lox*^ and *Pdgfrα*^*CreERT(+/-)*^*;Gcamp5-tdTomato*^*Lox/Lox*^ transgenic mice (Fig. 1a). Ca^2+^ imaging in acute slices was performed in LPC-induced lesions of corpus callosum as these lesions have several advantages: 1) since these are focal lesions, they are localized in specific regions, 2) they can easily be identified by DIC video microscopy at a low magnification as a brighter area than the surrounding non-lesioned dark-appearing white matter (Sahel et al., 2015) and imaging can then be performed at a high magnification inside the core of the lesion by either one- or two-photon Ca^2+^ imaging (Fig. 1b, 1d, 1e). In the LPC model, the response of oligodendroglia to demyelination occurs in a stepwise manner (migration of OPCs to the lesion site, OPC proliferation, OPC differentiation and remyelination), such that Ca^2+^ signals of oligodendroglia at each step of the remyelination process can be analyzed separately (Sahel et al., 2015). Tamoxifen injections to induce Cre expression a few days before the stereotaxic LPC injection are sufficient to elicit a good level of GCaMP expression in oligodendroglia 7 days post-injection (dpi) of LPC (Fig. 1b, 1d, 1e). This protocol led to a lesion-specific expression of GCaMP, as none was observed in intact corpus callosum, probably because adult PDGFRα^+^ oligodendrocyte progenitors revert to an immature state and shorten their cell cycle upon demyelination (Supplementary Fig. 1; Moyon et al., 2015; Cayre et al., 2021). In this work, we imaged oligodendroglia in *ex vivo* demyelinated brain slices by two different techniques: one-photon microscopy used to study the whole oligodendroglial population’s Ca^2+^ activity in a wide field and conventional two-photon microscopy used to study the oligodendroglial Ca^2+^ signals at the single cell level (Fig. 1b, 1d, 1e).

**Figure 1.**
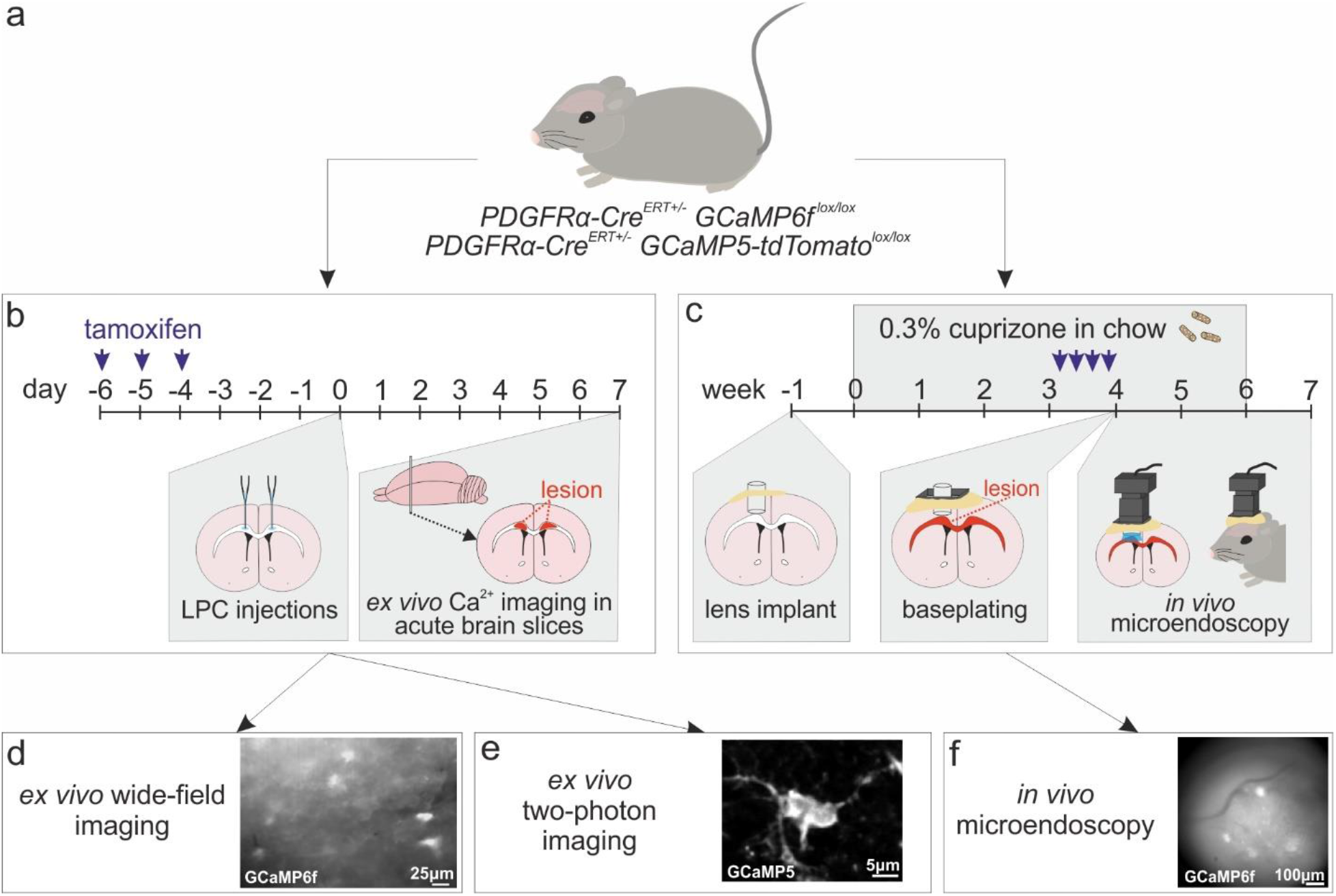
Experimental workflow for oligodendroglial Ca^2+^ imaging of *ex vivo* and *in vivo* preparations. **(a)** Demyelinating lesions were performed in adult *Pdgfrα*^*CreERT(+/-)*^*;Gcamp6f*^*Lox/Lox*^ and *Pdgfrα*^*CreERT(+/-)*^*;Gcamp5-tdTomato*^*lox/Lox*^ transgenic mice. **(b)** Four days after tamoxifen injection, demyelination was induced by LPC injection in the corpus callosum and *ex vivo* brain slices were performed 7 days after LPC injection. **(c)** Demyelination was induced by CPZ ingestion starting 1 week after GRIN lens implantation and tamoxifen injections were performed during the fourth week of CPZ ingestion. *In vivo* microendoscopy was performed from the fifth week of CPZ feeding and during the first week after CPZ withdrawal. **(d-f)** Ca^2+^ imaging in lesions was performed using wide-field microscopy (d) and two-photon microscopy (e) in *ex vivo* brain slices and wide-field microendoscopy (f) in freely moving mice.

After successfully imaging the Ca^2+^ dynamics in demyelinated lesions in *ex vivo* brain slices, we extended our approach by including *in vivo* Ca^2+^ imaging in demyelinated lesions. The main difficulty encountered when imaging the mouse corpus callosum *in vivo* is related to the depth of this region, particularly at the level of the motor cortex where demyelinated lesions are often analyzed (Ortiz et al., 2019). Although some lesions occur in the grey matter in MS, most of them actually occur in the white matter; therefore, performing oligodendroglial Ca^2+^ imaging in corpus callosum lesions is particularly relevant for this disease. For this purpose, we used *in vivo* one-photon microendoscopy which allowed us to visualize Ca^2+^ signals of oligodendroglia in the corpus callosum, at 1.8 mm from the surface of the brain. In order to monitor oligodendroglial Ca^2+^ signaling in callosal demyelinated lesions of freely moving mice (Fig. 1c), we operated the open-source Miniscope V4 device, often used to follow neuronal Ca^2+^ activity (Shuman et al., 2020; Cai et al., 2016). One interesting challenge in our approach was the fact that, although the LPC-induced demyelination model is appropriate for brain slices, demyelination is already obtained at 3 dpi (Sahel et al., 2015) and thus incompatible with *in vivo* microendoscopy which requires a multi-step procedure extending over several weeks (Fig. 1c). We therefore used the CPZ demyelination model in which intense demyelination occurs from the fourth week after CPZ ingestion without any surgical intervention (Fig. 1c; Remaud et al., 2017). For these experiments, a GRIN lens was implanted deep in the cortex above the target imaging region seven days before the start of the CPZ treatment. Five weeks after the GRIN lens implantation, a baseplate was mounted onto the animal’s skull providing fixtures to attach the Miniscope and allowing us to set the working distance between the objective and the lens (Fig. 1c). The tissue can thus recover for five weeks and any inflammation or blood accumulation under the lens can be cleared before the beginning of Ca^2+^ imaging experiments (Zhang et al., 2019). Tamoxifen was injected during the fourth week of CPZ treatment, when demyelination is advanced. Finally, *in vivo* imaging was done during demyelination to record Ca^2+^ signals from oligodendroglia in the demyelinated corpus callosum of freely moving mice (Fig. 1c, 1f).

### Occam and post-prOccam: Ca^2+^-signal analysis softwares for investigation of oligodendroglia in demyelinated lesions

To perform an automated and unbiased detection of active regions in different Ca^2+^ imaging datasets from demyelinated preparations, we developed a configurable Fiji/ImageJ2 plugin named Occam (Oligodendroglial cells calcium activity monitoring; Fig. 2a). We first validated our analytical workflow on wide-field one-photon Ca^2+^ imaging data recorded in acute slices of corpus callosum which revealed high levels of spontaneous Ca^2+^ activity in oligodendroglia of LPC-induced demyelinated lesions (Fig. 1b, 1d, 2a, 3a; Supplementary Video 1). Occam first performs noise and bleaching corrections of image stacks (Fig. 2a) and then uses the trainable machine learning-based WEKA Fiji/ImageJ2 segmentation plugin (Arganda-Carreras et al., 2017) combined with a local maxima segmentation tool to define regions of interest (ROIs) with fluctuating Ca^2+^ signals with high and medium pixel intensities (Supplementary Manual). Several parameters of ROI detection can be configured by the user to optimize Occam’s performance and to obtain an automated and unbiased ROI designation (for instance, the minimum size of an active ROI or a bleaching correction option). Once the ROI designation process is completed, Occam saves two files: one file stores each ROI area size in pixels and the coordinates of the geometric center of the ROI, the other file describes each ROI’s fluorescence behavior over time as a vector listing the mean fluorescence pixel intensity value in each frame contained in the image stack. The ROI fluorescence intensity vector is also referred to as the ROI trace (Supplementary Fig. 2a, 2b; Supplementary Manual). These two files are then read by the post-prOccam software to perform a systematic ROI analysis along with quantitative calculations ultimately aimed at easing the detailed characterization of oligodendroglial Ca^2+^ activity (Fig. 2b). The post-prOccam software was implemented as a configurable program where users can set various analysis parameters in a configuration file so as to adapt its operation to any specific dataset (Supplementary Manual).

**Figure 2.**
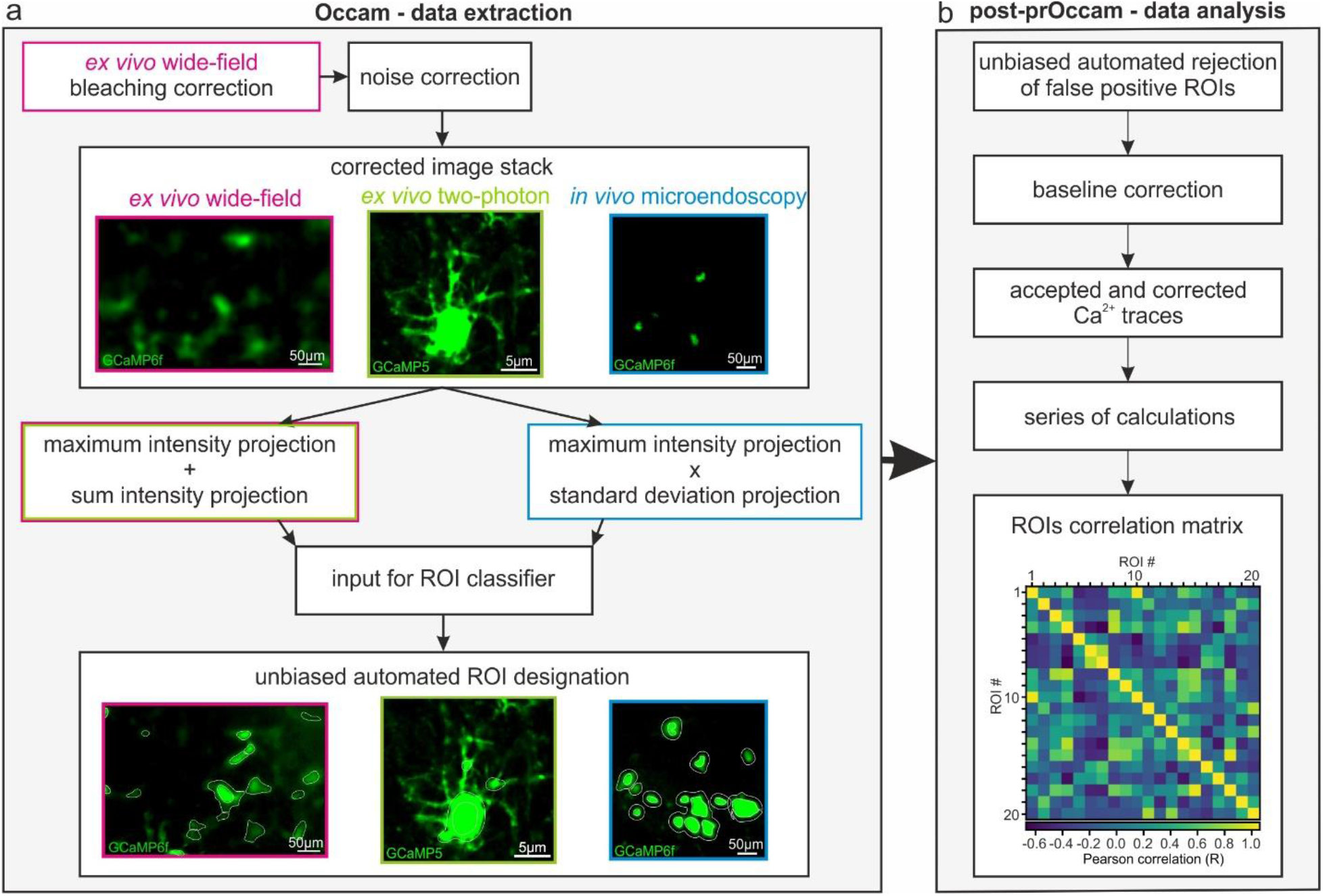
Occam and post-prOccam: an open source, fully automated and configurable analysis toolkit for oligodendroglial Ca^2+^ imaging of *ex vivo* and *in vivo* preparations. **(a)** The Occam software is available as a Fiji (tested in version 1.53t and earlier) plugin and configurable for the analysis of wide-field, two-photon and *in vivo* microendoscopy Ca^2+^ imaging. Occam performs bleaching correction only on *ex vivo* wide-field image stacks and does noise correction according to imaging condition (Supplementary Manual). Then, it uses the maximum and sum intensity projections for *ex vivo* image stacks and the maximum and standard deviation projections for *in vivo* image stacks to build an input for the ROI classifier. The ROI classifier of Occam defines ROIs with significant Ca^2+^ fluctuations fully automated and is used by WEKA for ROI designation. **(b)** Output of Occam can be fed to post-prOccam which is a configurable Python-based software that first rejects any ROIs that do not show significant Ca^2+^ fluctuations, then performs baseline corrections, subsequently performs a series of calculations for each ROI in the image stack and finally computes a ROIs correlation matrix of the Pearson correlation value of each ROI as compared to every other ROI. Occam and post-prOccam are available as free, multiplatform and open source softwares at: *https://gitlab.com/d5674/occam* and procedures are described in more detail in the Supplementary Manual.

**Figure 3.**
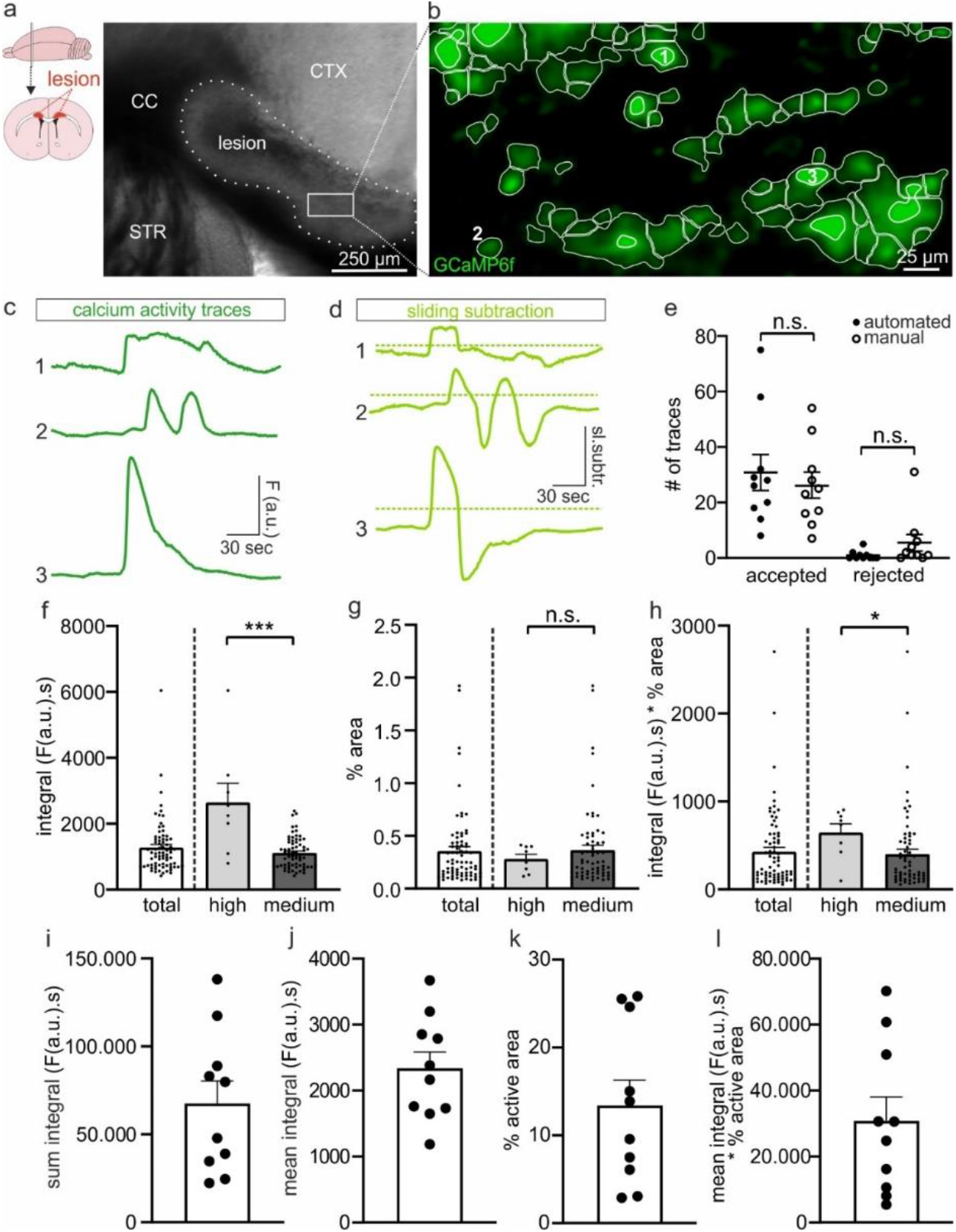
Using Occam and post-prOccam for analysis of ex vivo wide-field oligodendroglial Ca^2+^ imaging from a callosal LPC-induced demyelinated lesion. **(a and b)** Representative image of a demyelinated lesion in 4x DIC microscopy (a) and of GCaMP6f expressing oligodendroglia in 40x wide-field fluorescence imaging displaying detected active ROIs (white) as obtained with Occam (b). **(c and d)** Representative corrected ROI traces (c) and sliding window subtraction traces (d) as obtained with post-prOccam. **(e)** Comparison of the manual rejection of false positive ROIs and the automatic rejection of false positive ROIs by post-prOccam revealed no differences in accepted and rejected ROIs, validating post-prOccam’s performance. n.s.: not significant, two-way ANOVA followed by a Bonferroni multiple comparisons test. **(f-h)** Calculations performed by post-prOccam on each individual ROI of the image stack include amongst others the integral (f), percentage of active area (g) and integral multiplied by percentage of active area (h) and are separated for high and medium intensity ROIs. *p<0.05 and ***p<0.001; Mann-Whitney test. **(i-k)** Calculations performed by post-prOccam on all ROIs in an image stack include amongst others the sum integral (i), the mean integral (j), the percentage active area (k) and the mean integral multiplied by the percentage active area (l) (n=10 stacks, n=10 slices, n=7 mice). Error bars represent standard error of the mean. Dot plots are presented as mean±s.e.m.

Automated ROI detection algorithms cannot be error-free and almost always require an *a posteriori* rejection of false positive ROIs (Cantu et al., 2020). We developed the post-prOccam program that automatically fetches the data written by Occam and proceeds to an automatic rejection of ROIs lacking significant Ca^2+^ signal fluctuations. To reject false positive ROIs, we adapted an algorithm previously used for the fluorescence-based tracking of exocytotic events (Yuan et al., 2015; Supplementary Manual). Based on repeated intensity values subtraction in a sliding window over the whole ROI vector of mean intensity values (Supplementary Fig. 2a-c), our method allowed us to single out fluorescence intensity changes in an image stack that were greater than noise fluctuations (Fig. 2b, 3a-d). Since Ca^2+^ signals of oligodendroglia in lesions were characterized by long-lasting kinetics (Fig. 3c; rise time: 54.63±9.67 frames, equivalent to 35.25±5.36 s, n=34 events from n=8 fields), the sliding window subtractions were computed between points distanced by 40 frames, as: intensity = intensity(ROI[n+40]) - intensity(ROI[n]), with n being the frame number in the stack (Supplementary Fig. 2b, 2c). Each initial ROI trace is thus replaced by a new one, as computed from the sliding window subtractions, that is then tested against user-defined threshold parameters set in the configuration file (Fig. 3d; Supplementary Files 1-3). These threshold settings configure the rejection of ROI traces depending on both their Ca^2+^ event kinetics and noise. To find the best settings to eliminate false positive traces, we empirically changed the threshold parameters and compared the results with those obtained by a manual rejection of ROIs in several imaging stacks (Fig. 3e). The threshold values that were found to be the most effective in rejecting false positive ROIs were set as default values in the configuration file and provided in Supplementary File 1 (Fig. 3e; see detailed description of the threshold parameters in the Supplementary manual).

Once the accepted ROIs have been selected, post-prOccam performs a baseline subtraction for each ROI trace by subtracting a mean minimum intensity value previously calculated from a configurable number of trace points (Supplementary File 1; Supplementary Manual). We used a baseline subtraction computation rather than the conventional [(F(t)-F_0_)/F_0_] computation because we adapted the analysis to the Ca^2+^ dynamics of oligodendroglia in lesions. Indeed, the oligodendroglial cells studied in this report exhibited high overall levels of spontaneous Ca^2+^ activity, often right at the start of the recording, making it difficult to determine with certainty the F_0_ value of the resting fluorescence intensity. Finally, the post-prOccam program performs a series of calculations to help characterize the Ca^2+^ activity of oligodendroglia in lesions (Fig. 2b). Figure 3f-h illustrates part of the calculations performed for each ROI in a single image stack, *i*.*e*. the mean integral, the percentage of active area and the mean integral multiplied by the percentage of the active area. Since the complex features of oligodendroglial Ca^2+^ events preclude their proper individual isolation (Supplementary Fig. 3), we found it more appropriate to calculate the mean intensity integral of each ROI trace rather than to use a procedure for single event detection (see discussion). As expected, high intensity ROIs exhibited significantly larger mean integrals than medium intensity ROIs (Fig. 3f) and, despite similar mean percentage of active area between high and medium intensity ROIs, the mean integral multiplied by the percentage of the active area remained larger for high intensity ROIs (Fig. 3g-h). However, when considering all the analyzed stacks, the number of high intensity ROIs was always largely smaller than the number of medium intensity ROIs (3.5±0.9 high intensity ROIs, n=27.3±5.7 medium intensity ROIs and n=30.8±6.5 total ROIs for n=10 stacks). Moreover, the mean integral of medium intensity ROIs and that of all pooled data (Total) remain similar (Fig. 3f). Therefore, we hereinafter pooled the data from high and medium intensity ROIs for quantifications, but kept their detection separate because the WEKA plugin used by Occam performed better when classifying ROIs in these two categories. The average calculations for all ROIs in several image stacks show that Ca^2+^ activity between demyelinated lesions at 7 dpi can be variable but sufficiently high to always allow us to detect changes over only a few minutes of recording (Fig. 3i-l).

For informational and debugging purposes, a log file is produced by post-prOccam with a highly detailed account of all the data processing steps and their outcome. The accepted ROIs and the corresponding processed data of a stack are saved by post-prOccam in two separate files for further statistical analysis. Of note is the pp-supervisor program that allows the scientist to run post-prOccam automatically over a set of stacks located in a single directory using the configuration file that was elaborated during the exploratory steps described above.

### Interval-based analysis and evaluation of synchronized Ca^2+^ signals of oligodendroglia in lesions

One important feature of post-prOccam is its ability to compare ROIs only at specific time point intervals of the recorded datasets. For instance, in the case of applications of pharmacological agents during Ca^2+^ imaging recordings, a user might wish to compare Ca^2+^ signals in the absence or in the presence of a drug in a single image stack. The default post-prOccam software behavior is to perform all the calculations described in the previous sections on the whole ROI vector. However, in specific cases mentioned above, it might be useful for the calculations to be performed only over selected ranges of the ROI vector (that is, intervals of that ROI’s acquisition time points). The configuration file provides a section in which the user might list any number of ROI vector intervals over which to perform the previously described calculations. To validate this feature, we bath-applied the muscarinic receptor agonist carbachol in the presence of a cocktail of antagonists to stimulate intracellular Ca^2+^ signals of oligodendroglia during one-photon Ca^2+^ imaging recordings of 7 dpi callosal LPC-induced lesions in brain slices (Fig. 4a, 4b). Oligodendroglia express muscarinic receptors M1, M3 and M4, which, when activated by carbachol, increase intracellular Ca^2+^ signals (Abiraman et al., 2015; Cohen & Almazan, 1994; Welliver et al., 2018). In order to determine the extent of the Ca^2+^ signal activity changes upon the bath application of carbachol, we directed post-prOccam to compare the Ca^2+^ signals in each ROI only during two distinct acquisition intervals: the control interval before the carbachol application and the interval during which the carbachol was applied. As expected, we observed that a bath application of 50 μM carbachol in the presence of a cocktail of antagonists indeed induced an increase in intracellular Ca^2+^ signals in oligodendroglia, as revealed by an increase in both the mean integral and the mean integral multiplied by the percentage of the active area, compared to the control (Fig. 4a-d).

**Figure 4.**
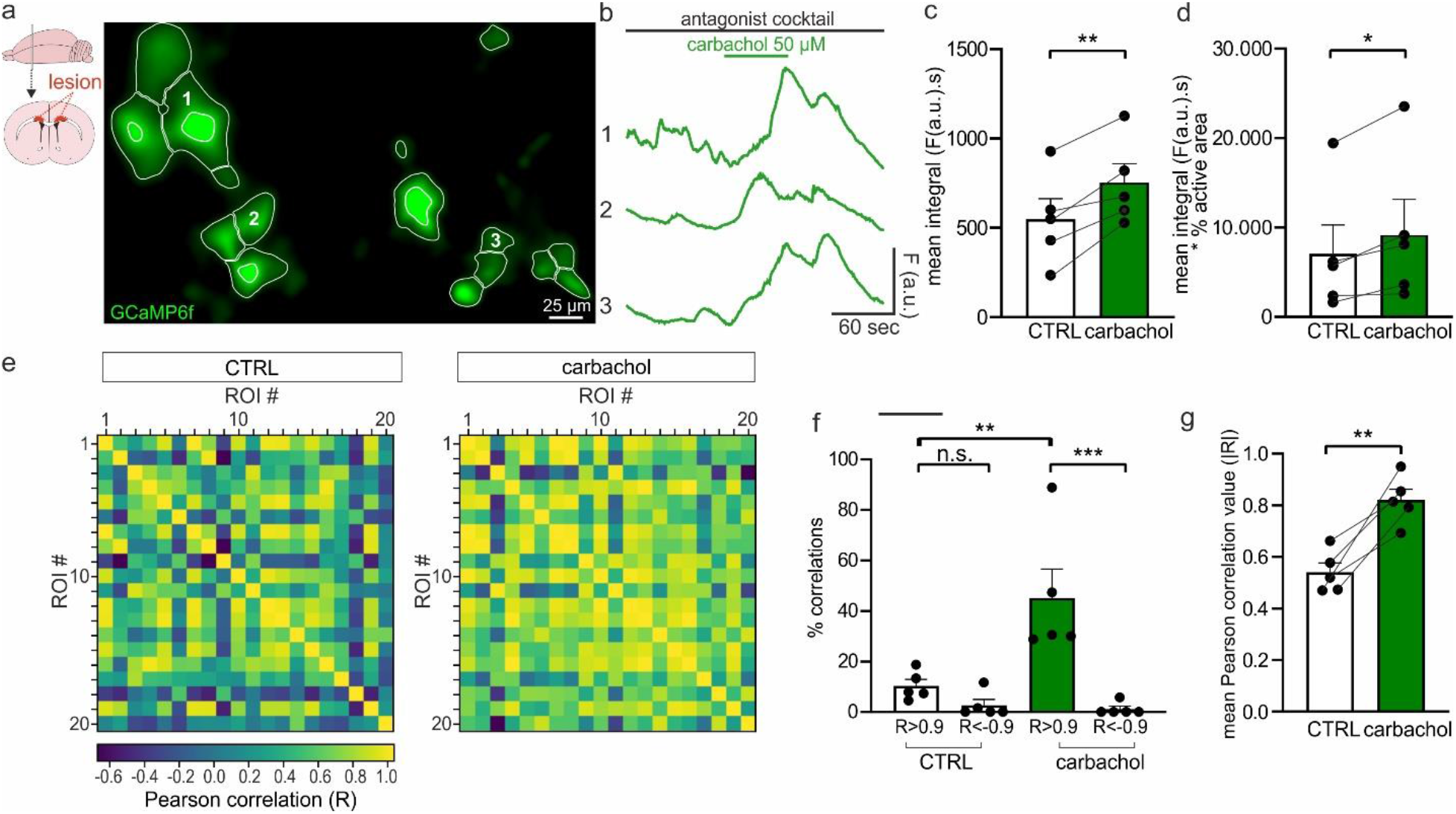
Using Occam and post-prOccam for interval analysis and evaluation of synchronized Ca^2+^ signals of oligodendroglia in LPC-induced demyelinated lesions. **(a)** Representative image of Ca^2+^ imaging in callosal LPC-induced lesions in *ex vivo* brain slices that were exposed to 50 μM carbachol to induce increases in Ca^2+^ signals in oligodendroglia in the presence of an antagonist cocktail containing 10 μM NBQX, 50 μM AP5, 10 μM GABAzine, 1 μM TTX and 50 μM mecamylamine. The image displays detected active ROIs (white) as obtained with Occam. **(b)** Representative Ca^2+^ traces showing Ca^2+^ increases during carbachol application as obtained with post-prOccam. **(c and d)** Mean integral (c) and mean integral multiplied by the percentage of active area (d) of Ca^2+^ signals in control and after exposure to carbachol. *p<0.05, **p<0.01; paired Student’s t-test. **(e)** Example of correlation matrixes obtained with post-prOccam before and after carbachol exposure. Each square indicates the Pearson correlation value of one ROI with another. Yellow indicates high positive Pearson correlation, while dark blue indicates high negative correlations. Note that traces 1, 2, 3 in b correspond to ROIs 7, 12, 14 in the matrix. **(f)** The percentage of correlations in the correlation matrix that is significantly negative (R<-0.9) or significantly positive (R>0.9) in both control and carbachol exposed conditions. n.s.: not significant, **p<0.01, ***p<0.001, two-way ANOVA followed by a Tukey’s multiple comparisons test. **(g)** Mean Pearson correlation values for control and carbachol conditions. CTRL: control before carbachol exposure (n=5 stacks, n=5 slices, n=4 mice). **p<0.01; paired Student’s t-test. Dot plots are presented as mean±s.e.m.

While it is now established that a population of astrocytes may exhibit high levels of synchronized Ca^2+^ activity *in vitro* and *in vivo* (Kolzumi et al., 2010; Ingiosi et al., 2020), nothing is known about the potential of Ca^2+^ signals in oligodendroglia to be synchronized. We thus tested this possibility in demyelinating lesions by implementing a correlation calculation which allows the user to establish whether ROIs within a given image stack show synchronized Ca^2+^ signals. The post-prOccam program computes an inter-ROI Pearson correlation coefficient matrix between each ROI trace and every other ROI trace (Fig. 4e; see Materials and Methods; Ingiosi et al., 2020). In control conditions, we found that most ROIs did not show any correlated Ca^2+^ activity, as evidenced by a low percentage of correlated ROIs (|R| < 0.9) and a mean Pearson correlation coefficient R value of 0.54±0.24 (Fig. 4f-g). The remaining few ROIs that displayed correlated Ca^2+^ activity had mainly a positive rather than a negative correlation (Fig. 4f). Interestingly, upon application of the muscarinic agonist carbachol in the presence of a cocktail of antagonists, a significant overall increase of the positive inter-ROI Ca^2+^ activity correlation was observed (Fig. 4e-g), with a mean Pearson correlation value of 0.82±0.04. These data showed that the majority of ROIs did simultaneously respond to the agonist. The post-prOccam software thus successfully determined the degree of Ca^2+^ activity correlation between ROIs and was able to detect simultaneous increases of that activity as induced by pharmacological agents. In order to adapt to any specifics of biological applications, the Pearson correlation R value threshold might be configured. The percentage of correlated ROIs as well as the mean Pearson correlation coefficient are reported for each image stack in the corresponding output file generated by the program. Taken together, our results show that the Occam and the post-prOccam software programs make for a configurable and automatable solution for the analysis of oligodendroglial Ca^2+^ signals.

### Two-photon imaging of Ca^2+^ signals in single oligodendroglia in demyelinated lesions

Although one-photon imaging allows one to determine the global Ca^2+^ signaling properties of oligodendroglia in a large area of the LPC-induced lesion, the spatial resolution of this technique is not sufficient to reveal local Ca^2+^ changes occurring specifically either in the soma or in the cell processes of any individual cell. To test whether our workflow is suitable for the analysis of images obtained by two-photon microscopy at the single cell resolution, we performed two-photon Ca^2+^ imaging of putative OPCs and OLs in callosal LPC-induced lesions in *ex vivo* brain slices and analyzed the data using Occam and post-prOccam (Fig. 5; Supplementary Video 2 and Supplementary File 2). We used morphological criteria to distinguish OPCs and OLs, and only recorded cells for which we could connect the processes to a particular soma. In corpus callosum, OPCs were characterized by a relatively small round soma and a stellate arborization with thin processes (Fig. 5a; Chittajallu et al., 2004) whereas OLs had a larger soma and principal processes often aligned with axons (Fig. 5f; Bakiri et al., 2011). Morphological determination of putative OPCs and OLs allowed us to reveal that, even though the area of ROIs in the processes is significantly reduced as compared to that of ROIs in the soma, the mean intensity integral of their Ca^2+^ signals is similar to that of Ca^2+^ signals from the cell soma (Fig. 5a-c, 5f-h). Although the mean number of ROIs in OPCs and OLs was similar (9.44±1.97 in OPCs vs 8.00±2.19 in OLs) as was the rise time of Ca^2+^ events (3.61±1.14 s n=10 events from 5 fields for OPCs vs 4.16±0.68 s; n=37 events from 6 fields for OLs p=0.71 for soma and process data pooled together, unpaired Student’s t-test), Ca^2+^ signals had a higher integral in OPCs than in OLs (Fig. 5 a-c, 5f-h, p=0.0391 for soma and process data pooled together, unpaired Student’s t-test)). Altogether, these results not only show high levels of Ca^2+^ activity in processes of OPCs and OLs, but also that this activity is greater in OPCs inside demyelinating lesions. Importantly, we also observed that, whatever the cell type involved (OPCs or OLs), the Ca^2+^ signals were poorly correlated, which suggests that they occurred independently in the soma and in the processes and at different subcellular locations (Fig. 5 d-e, 5i-j). In summary, these results show that Occam and post-prOccam can successfully determine the characteristics of oligodendroglial Ca^2+^ signals recorded by different imaging techniques and at different spatial scales.

**Figure 5.**
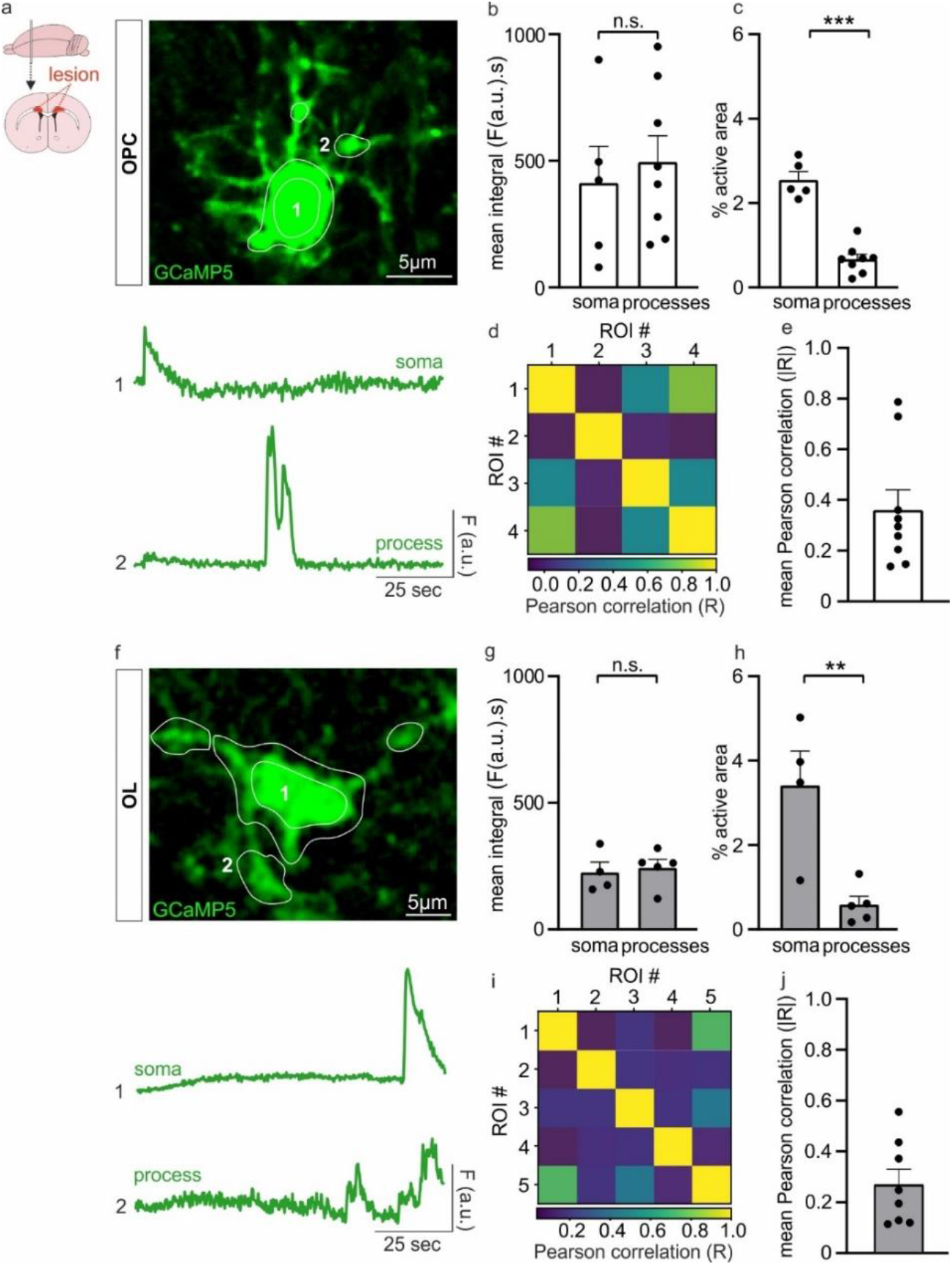
Using Occam and post-prOccam for the analysis of two-photon oligodendroglial Ca^2+^ imaging in callosal LPC-induced demyelinated lesions in *ex vivo* brain slices. **(a)** Representative image of a putative OPC **(a)** and a putative OL **(f)** expressing GCaMP5 with examples of corrected Ca^2+^ traces from the soma and a process (bottom) obtained with Occam and post-prOccam. **(b-c, g-h)** Mean integrals (b, g) and % active area (c, h) for the soma and the processes of OPCs (b-c) and OLs (g-h). n.s.: not significant. **p<0.01, ***p<0.001, unpaired Student’s t-test. **(d, i)** Examples of correlation matrices describing a lack of synchronization of Ca^2+^ signals in the soma and processes of OPCs (d) and OLs (i). Each square indicates the Pearson correlation value of one ROI with another ROI. Yellow indicates high positive Pearson correlation, while blue indicates no correlation. Note that traces 1 and 2 in a and f correspond to ROIs 1, and 2 in the matrices in d and i, respectively. **(e and j)** The mean correlation value of OPCs and OLs (n=9 stacks, n=9 slices, n=7 mice for OPCs and n=8 stacks, n=7 slices, n=6 mice for OLs). Dot plots are presented as mean±s.e.m.

### *In vivo* microendoscopy Ca^2+^ imaging of oligodendroglia during the demyelination process

Although *in vivo* Ca^2+^ imaging has been extensively studied in neurons and astrocytes, only few studies in the zebrafish have reported *in vivo* Ca^2+^ signals in oligodendroglia (Baraban et al., 2018; Krasnow et al., 2018; Marisca et al., 2020; Li et al., 2022). Moreover, no current information exists on oligodendroglial Ca^2+^ imaging in the demyelinated mouse brain *in vivo*. To fill this gap, we decided to adapt our experimental and analytical workflow to the study of oligodendroglial Ca^2+^ signals during demyelination in freely moving mice (Fig. 1c-f, Fig. 2a). As a proof-of-concept, we used the CPZ-induced demyelination model and performed microendoscopic Ca^2+^ imaging recordings of oligodendroglia in the corpus callosum of awake demyelinated mice (Fig. 6a-b; Supplementary Video 3). Four animals were implanted according to the procedure described in Figure 1. Three of them were recorded during the fifth postnatal week of CPZ feeding, when demyelination is advanced (Remaud et al., 2017), and sacrificed afterwards in order to confirm the proper position of the GRIN lens above the corpus callosum. The fourth mouse was maintained alive and imaged at four different timepoints to show that multiple recordings of the same animal can be made over several weeks.

**Figure 6.**
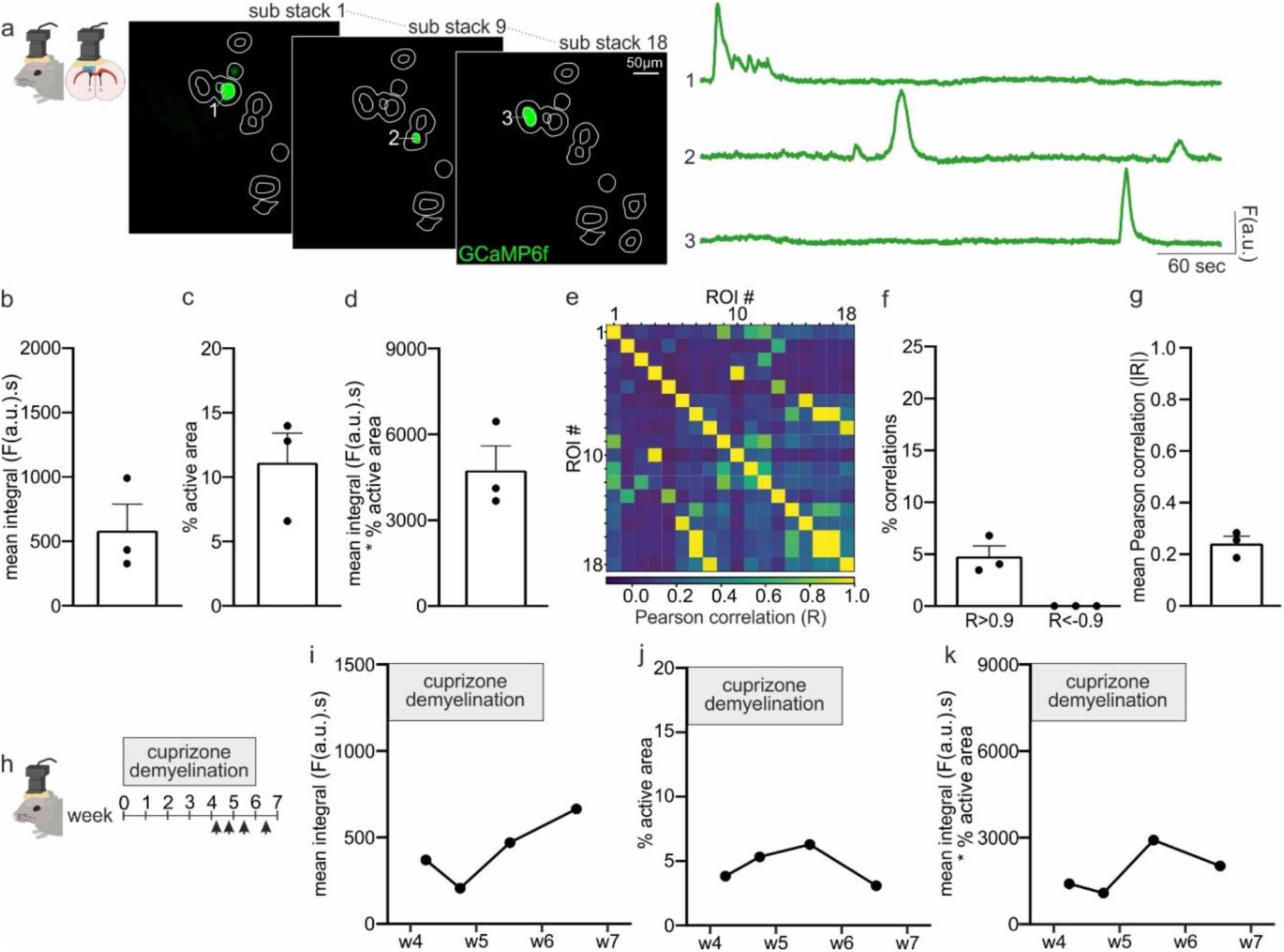
Using Occam and post-prOccam for the analysis of *in vivo* microendoscopy Ca^2+^ signals of oligodendroglia in the demyelinated corpus callosum of freely moving mice. **(a)** Representative images and corrected Ca^2+^ traces of microendoscopy Ca^2+^ imaging in freely moving mice during the fifth week of CPZ demyelination. The image displays detected active ROIs (white) in several sub-stacks as obtained with the *in vivo* analysis option of Occam. **(b-d)** Mean integral (b), % of active area (c) and mean integral multiplied by percentage of active area (d) during the fifth week of CPZ demyelination (n=3 mice). Data is presented as mean±s.e.m. **(e)** Correlation matrix of the *in vivo* experiment in a obtained with post-prOccam. Each square indicates the Pearson correlation value of one ROI with another ROI. Yellow indicates high positive Pearson correlation, while dark blue indicates no Pearson correlation. Note that trace 1, 2, 3 in a correspond to ROIs 1, 2, 3 in the matrix. **(f)** The percentage of correlations in the correlation matrix. **(g)** The mean Pearson correlation value. **(h)** *In vivo* microendoscopy Ca^2+^ imaging was performed in the same mouse at four timepoints, namely during weeks 5 and 6 of CPZ-induced demyelination and during week 7 after CPZ withdrawal. **(i)** Mean integral, **(j)** % of active area and **(k)** mean integral multiplied by percentage of active area during the fifth, sixth and seventh week in the same mouse.

The image stacks we acquired contained 4200 frames collected over 7 minutes. To enhance the precision of ROI detection by Occam in such large data files, the program first splits the image stack into a configurable number of sub-stacks. Then, for each sub-stack, it performs the noise correction, uses the maximum and standard deviation projections to build an input for the ROI classifier and proceeds to the automated ROI designation using the WEKA Fiji/ImageJ2 segmentation plugin combined with a local maxima segmentation tool (Fig. 2a; Supplementary Manual). This side-step from the *ex vivo* version of Occam allows for accurate detection of ROIs in large image stacks collected over many minutes because it also captures ROIs that are only briefly active and that would be overlooked when making a projection image over the entire imaging duration. However, this procedure can lead to an overestimation in the number of active ROIs if a specific region of the stack is active several times during a single experiment and, thus, is detected as a ROI in multiple sub-stacks. To overcome this problem, Occam projects all ROIs from all sub-stacks onto each other and allows the user to determine a degree of overlap for which overlapping ROIs are merged into a single ROI (Supplementary Manual). Like for *ex vivo* preparations, ROIs with high and medium pixel intensities were considered as displaying significant Ca^2+^ signal fluctuations. As the number of sub-stacks can be configured by the user, the ROI detection process will be successful even in recordings longer than 7 minutes, either in a single acquisition or in multiple acquisitions performed on the same animal over weeks (Fig. 6). In sum, Occam features for in vivo experiments have been adapted so that it can detect not only shortly occurring events, but also the repeated activation of a single region over a long experiment. Once the ROI selection process has been performed, the downstream processing by post-prOccam is identical to that described in the preceding sections (Supplementary File 3).

Our analysis revealed that oligodendroglia *in vivo* show high levels of spontaneous Ca^2+^ activity in the fifth week of CPZ-induced demyelination (Fig. 6a-d; mean number of ROIs of 15±4.58). We could identify ROIs with multiple Ca^2+^ events with complex and long-lasting kinetics of several seconds (Fig. 6a and Supplementary Fig. 3c). The mean rise time, however, was on average faster than in *ex vivo* wide-field experiments (rise time: 31.48±5.54 frames, equivalent to 6.29±0.91 s, n=48 events from n=3 fields). Furthermore, the low percentage of correlated ROIs and the small mean Pearson correlation coefficient showed that most ROIs did not display any correlated Ca^2+^ activity (Fig. 6e-g), confirming the lack of synchronized Ca^2+^ signals in oligodendroglia observed *ex vivo* in the LPC model. Finally, we performed *in vivo* microendoscopy Ca^2+^ imaging in the same mouse during weeks 5 and 6 of CPZ-induced demyelination and during week 7 after CPZ withdrawal (Fig. 6h). The mean pixel intensity integral, the percentage of active area and the mean integral multiplied by the percentage of the active area remained relatively stable during this period, indicating that the imaging conditions did not deteriorate over time (Fig. 6h-k). The monitoring of oligodendroglial Ca^2+^ activity over weeks in the same animal may give insight into the behaviour of this cell population during demyelination. To our knowledge, this is the first time that Ca^2+^ signals of oligodendroglia during demyelination have been recorded in real time and the successful validation of our workflow prompts future research on oligodendroglial Ca^2+^ signaling in the living injured mammalian brain.

## Discussion

While it is suspected that oligodendroglial Ca^2+^ signaling plays a key role in myelin repair, the characteristics of oligodendroglial Ca^2+^ signals in demyelinated lesions remain unknown. This is due to a limited number of *ex vivo* or *in vivo* Ca^2+^ imaging studies as well as a lack of automated analytical tools adapted to the monitoring of the specific characteristics of oligodendroglial Ca^2+^ activity. To fill this need, we devised experimental conditions for *ex vivo* and *in vivo* Ca^2+^ imaging in mouse demyelinated lesions and implemented an analytical workflow comprising two programs, Occam and post-prOccam, which provide a reliable solution for the automated in-depth analysis of one- and two-photon oligodendroglial Ca^2+^ imaging data. Tested in Windows 10 and Debian GNU/Linux, these cross-platform programs were designed to be easily configurable to ensure an unbiased selection of active ROIs and to allow an efficient analysis of large datasets. Licensed under the GNU GPLv3+ license, both programs can be freely used, modified according to any specific need and question, and redistributed. Since the Ca^2+^ signal detection is finely configurable in both programs, the use of our software solution might be extended to new use cases such as other glial cell types.

By harnessing the features in these two programs, we could monitor the spontaneous oligodendroglial Ca^2+^ activity in different toxic models of demyelination, both *ex vivo* and *in vivo*. In any of these possible configurations, Ca^2+^ traces that were first obtained with Occam then undergo a post-processing procedure with post-prOccam that processes them to reject ROIs with too small Ca^2+^ fluctuations (false positives), which can be construed as a refinement of the ROI detection process. We found that this automatic Ca^2+^ trace selection in our workflow performs equally well as a manual Ca^2+^ trace selection, thus validating the procedure. Among other analysis options, the ability to restrict the processing and quantification steps to specific acquisition intervals has proven useful to screen the effect of drugs on oligodendroglial Ca^2+^ signaling. Furthermore, a number of features in post-prOccam make it possible to spot eventual synchronized Ca^2+^ signals across different ROIs in a given frame stack. Here we successfully applied these analysis modalities to detect, quantify and correlate Ca^2+^ activity increases upon carbachol bath application to demyelinated brain slices in order to activate muscarinic receptors in oligodendroglia, as previously described in cell culture (Cohen & Almazan, 1994; Welliver et al., 2018). Interestingly, muscarinic receptor agonists can increase OPC proliferation, but block OPC differentiation and remyelination (De Angelis et al., 2012; Abiraman et al., 2015; Welliver et al., 2018). Moreover, the muscarinic receptor antagonist benztropine was found to be beneficial to induce myelin repair in an animal model (Deshmukh et al., 2013) and the muscarinic receptor antagonist clemastine was shown to improve myelin regeneration in a clinical trial with multiple sclerosis patients (Green et al., 2017). The positive effect of these antagonists might thus be caused by the blockade of muscarinic receptor-dependent Ca^2+^ signaling mechanisms, which is a subject for further research. These studies highlight the importance of investigating the Ca^2+^ response of oligodendroglia to novel remyelinating drugs. The workflow described in this report will facilitate these future studies.

Oligodendroglial Ca^2+^ events are unique in their variability and duration and are thus better qualified using the ROI-based measurements number, size and mean pixel intensity integral instead of the conventional measurements such as frequency, amplitude and duration of isolated events. Indeed, unlike neurons, that display well-defined Ca^2+^ signals on a millisecond scale (Chua & Morrison, 2016), oligodendroglia show complex Ca^2+^ events characterized by very slow and variable kinetics which make their detection and isolation difficult, in particular because of their frequent convolution (Supplementary Fig. 3a-c). This complexity of oligodendroglial Ca^2+^ events has previously been observed with two-photon microscopy (Balia et al., 2017) and in recordings at single cell resolution (Baraban et al., 2018; Krasnow et al., 2018; Battefeld et al., 2019; Marisca et al., 2020), indicating that complex Ca^2+^ dynamics are a hallmark of oligodendroglia and it is therefore essential to evaluate them in an appropriate manner. Thresholding techniques or event template detection methods commonly used on neuron and astrocyte Ca^2+^ imaging datasets are not easily applicable to the unique and complex Ca^2+^ events observed in oligodendroglia. To overcome this constraint, we evaluate activity levels by measuring the number and size of active ROIs as well as the integral of the traces. Our measurements thus account for the activity throughout the whole ROI trace or during configurable intervals without isolating Ca^2+^ activity events. In the eventuality that a detailed description of Ca^2+^ events would be desirable, extra measurements may be performed either manually or by other post-processing programs on the corrected active ROI traces as output by post-prOccam. In our case, we manually inspected these corrected traces and could establish that the mean rise time of Ca^2+^ events easily isolated by eye lasted from few seconds to minutes, independently of the imaging condition (Supplementary Fig. 3). These slow kinetics are in line with the half-width duration of 9 s reported for myelin internodes in the neocortex (Battefeld et al., 2019), but our results also show that oligodendroglial Ca^2+^ events in lesions are poorly correlated and may have long variable durations.

We could successfully and automatically detect oligodendroglial-related Ca^2+^-active ROIs in wide field as well as in single cell Ca^2+^ imaging frame stacks obtained in brain slices. These *ex vivo* recordings revealed significant but uncorrelated spontaneous Ca^2+^ activity in oligodendroglia inside demyelinated lesions, both at the population level and at the individual cell level. In particular, our two-photon Ca^2+^ imaging experiments showed that Ca^2+^ signals in ROIs located in soma or processes are not correlated, suggesting that they occur independently in subcellular compartments. In this configuration, our results also indicate that, compared to OLs, OPCs displayed a higher Ca^2+^ activity whose significance remain to be elucidated. The experimental and analytical tools described in this study will help to further explore the characteristics of these signals at different states of cell maturation and at different time points of the demyelination and remyelination process. Importantly, our workflow for acquiring and analyzing *in vivo* oligodendroglial Ca^2+^ signals will also allow users to investigate the role of these signals in the mammalian brain in real time. We were able to corroborate that the oligodendroglial Ca^2+^ activity observed *in vivo* is compatible with that observed in brain slices because it was variable but significant, poorly correlated and characterized by complex slow kinetics. We also showed that Ca^2+^ activity can be stable in the same mouse from one week to another. This type of *in vivo* experiments opens up new perspectives for analyzing how the oligodendroglial Ca^2+^ activity changes over time during myelin repair and when the mouse is performing a behavioral task.

In conclusion, our *ex vivo* and *in vivo* results suggest that oligodendroglial Ca^2+^ signaling plays an important role during the demyelination process. Therefore, the presented experimental and analytical framework will aid future investigations into oligodendroglial Ca^2+^ signaling during demyelination and myelin repair. As such, it might contribute to the elucidation of Ca^2+^-related mechanisms implicated in the success or failure of remyelination in demyelinating diseases such as MS.

## Materials and Methods

### Experimental animals

All experiments followed European Union and institutional guidelines for the care and use of laboratory animals and were approved by both the French ethical committee for animal care of the University Paris Cité (Paris, France) and the Ministry of National Education and Research (Authorization N° 13093-2017081713462292). They were performed with male and female *Pdgfrα*^*CreERT(+/-)*^*;Gcamp6f*^*Lox/Lox*^ *or Pdgfrα*^*CreERT(+/-)*^*;Gcamp5-tdTomato*^*Lox/Lox*^ transgenic adult mice (7 to 9 weeks old) obtained by crossing *Pdgfrα*^*CreERT*^ (stock 018280, The Jackson Laboratory) with *Ai95(Rcl-Gcamp6f)-D* (stock 028865, The Jackson Laboratory, USA) or *Gcamp5-tdTomato*^*Lox/Lox*^ (stock 028865, The Jackson Laboratory, USA). Animals were genotyped by PCR using specific primers for Cre. These mice express GCaMP6f or GCaMP5 in OPCs and their progeny upon tamoxifen injection. Tamoxifen (1 mg in miglyol oil, Caesar & Loretz, Germany) was administered intraperitoneally once a day for three consecutive days starting seven days before inducing demyelination by LPC injection or during week 4 of CPZ administration (Fig. 1a-c). All animals had *ad libitum* access to food and water and were exposed to a 12 hr light/dark cycle, a controlled average temperature of 21 °C and 45% humidity.

### Intracranial injections of LPC to induce demyelinated lesions

Mice were deeply anaesthetized using ketamine/xylazine (100/5 mg/kg, i.p.) or isoflurane (1.5 %) and headfixed in a stereotaxic frame (Kopf Instruments, USA). Before starting the surgery, mice received a subcutaneous injection of buprenorphine (0.1 mg/kg) as analgesic. Throughout surgical procedures mice were placed on a 37 °C heating blanket and ophthalmic dexpanthenol gel (Chauvin Bausch & Lomb GmbH, France) was applied to the eyes to prevent dehydration. The skin on the head was treated with betadine and lidocaine and opened to expose the skull. The head was aligned using both lambda and bregma, and two 0.50 mm diameter holes were bilaterally drilled in the skull at coordinates +1.40 mm anterior from bregma and +0.95 and –0.95 mm lateral from the midline. The dura was removed and a glass pipette connected to a Hamilton syringe containing LPC (1% in PBS; Merck, Germany) was lowered into the brain until 1.80 mm depth from the brain surface reaching into corpus callosum. The pipette rested in the brain for four minutes and subsequently 0.8 μL of LPC was injected two times with 4 minutes between injections. After another 4 minutes the pipette was slowly retracted from the brain and the LPC injection procedure was repeated in the other hemisphere. Skin was sutured (Mersilene®, EH7147H; Ethicon, USA) and cleaned with betadine and mice were allowed to recover at 37 °C before returning to their home cage.

### Acute brain slice preparation

Acute coronal slices of 300 μm containing demyelinated lesions were prepared as previously described (Mozafari et al., 2020). Brain slices were cut using a vibratome (Microm HM 650V, Thermo Scientific, USA) in a chilled cutting solution containing (in mM): 93 NMDG, 2.5 KCl, 1.2 NaH2PO4, 30 NaHCO3, 20 HEPES, 25 Glucose, 2 urea, 5 Na-ascorbate, 3 Na-pyruvate, 0.5 CaCl2, 10 MgCl2 (pH to 7.3-7.4; 95% O2, 5% CO2) and kept in cutting solution for 6 to 8 min at 34°C. Slices were transferred to a standard extracellular solution at 34°C for about 20 min. Extracellular solution contained (in mM): 126 NaCl, 2.5 KCl, 1.25 NaH2PO4, 26 NaHCO3, 20 Glucose, 5 Na-pyruvate, 2 CaCl2, 1 MgCl2 (pH to 7.3-7.4; 95% O2, 5% CO2).

### *Ex vivo* wide-field calcium imaging

Demyelinated lesions were recognized under the microscope at 4x as a brighter area in corpus callosum as described previously (Sahel et al., 2015). After identifying the demyelinated lesion at 4x, cells expressing GCaMP6f in the lesion were visualized with a 40x water immersion objective in a wide-field microscope (Olympus BX51) using a LED system (CoolLED PE-2; Scientifica, UK) and a CCD camera (ImageQ, Optimos; Scientifica, UK). Excitation and emission wavelengths were 470 nm and 525 nm, respectively. The CCD camera and the LED system were controlled using a Digidata 1440A interface and Pclamp10.5 software (Molecular Devices, USA). The image stacks were acquired at a frame rate of 1.75 Hz with 50 ms light exposure for a total duration of 240 s using Micro-manager-1.4 plugin under Fiji (version 1.53k or later) (Schindelin et al., 2012). Ca^2+^ imaging during bath applications of 50 μM carbachol were performed after incubating the slices for five minutes with an antagonist cocktail containing 10 μM NBQX, 50 μM AP5, 10 μM GABAzine, 1 μM TTX and 50 μM mecamylamine.

### *Ex vivo* two-photon calcium imaging

Two-Photon Ca^2+^ imaging was performed using a two-photon laser scanning microscope (Otsu et al., 2014). A 40x water-immersion objective (Olympus40x LumPlanFL N 540x/0.8) in combination with a 900 nm excitation beam from a femtosecond Ti:Sapphire laser (10 W pump; Mira 900 Coherent, Santa Clara, CA) was used to image GCaMP5 in individual putative OPCs and OLs identified by their morphology. Cells with round soma and numerous processes extending towards various directions were designated OPCs, while cells with elongated soma’s containing thicker and more T-shaped processes were designated OLs. GCaMP5 was detected with Hamamatsu photon counting PMTs through an emission filter (HQ500/40, Chroma). Ca^2+^ signals of individual cells were imaged in high resolution at 3.60-6.27 frames per second during 99 seconds.

### Cuprizone treatment and *in vivo* microendoscopy calcium imaging

One week before starting the CPZ treatment, mice (P35-P42) were subjected to GRIN lens implantation surgery (Fig. 1c). Animals were prepared for craniotomy in a stereotaxic apparatus as described above, and a 1 mm diameter circular cranial window was drilled around a midpoint at coordinates +1.40 mm anterior from bregma and +0.95 mm lateral from the midline. Using a blunt 23 G needle attached to a vacuum pump, the cortex was aspirated in the middle of the cranial window from the top of the brain surface until 1.40 mm depth. A 1.0 mm diameter GRIN lens (Inscopix, USA) was then implanted at 1.60 mm depth, just above the corpus callosum at the motor cortex level. This placement allows imaging inside this white matter region at 1.80 mm depth as the working distance of the GRIN lenses used is ± 200 μm. The lens was held in place by super glue and dental cement (Unifast, Japan) and covered with kwik-sil sealant (World Precision Instruments, USA) until the baseplate surgery. Animals were allowed to recover at 37°C before being transferred to their home cage.

For the seven days following the lens implantation procedure, animals were fed cherry or bacon flavored nutragel food (Bio-Serv, France). Subsequently, we supplemented the nutragel food with 0.3% CPZ (Sigma) for the 6 following weeks to induce demyelination. During week 4 of the CPZ diet, 4 daily 1 mg tamoxifen injections were performed intraperitoneally to induce GCaMP6f expression in demyelinated areas of *Pdgfrα*^*CreERT(+/-)*^*;Gcamp6f*^*Lox/Lox*^ mice. During week 5 of the CPZ diet, animals were subjected to a baseplating surgical procedure (Fig. 1c). Animals were anaesthesized with ketamine/xylazine (100/5 mg/kg, i.p.) and the kwik-sil seal was removed from the lens. Animals were then headfixed in a stereotaxic apparatus, and a miniscope V4 (Open Ephys, Portugal) with a baseplate connected to it was held over the lens until a clear image of the brain surface under the lens was obtained. We then cemented the baseplate onto the head of the mouse. A Ca^2+^ imaging stack was obtained during the baseplating procedure to ensure that the miniscope could efficiently detect Ca^2+^ signals from the demyelinated corpus callosum. Mice were recovered at 37°C and returned to their homecage. As a proof of concept, *in vivo* Ca^2+^ imaging was performed from week 5 of CPZ-induced demyelination (Fig. 1c). Image stacks were obtained in freely moving mice with a sampling rate of 10 frames per second during 7 minutes.

### Occam and post-prOccam: an automated analytical workflow for oligodendroglial Ca^2+^ imaging data

The analytical processing consists of two parts performed by two distinct software pieces thoroughly described in the software user manual provided as supplementary information (Supplementary Manual). Briefly, the analysis is composed of two main steps: first, the image stack is processed by Occam, a Fiji/ImageJ2-based plugin written in Java (Fiji/Imagej2 version 1.53k or later); second, the data output by Occam is processed by post-prOccam, a software written in Python, to both refine it and perform a series of calculations on the refined data. Importantly, *ex vivo* one-photon, *ex vivo* two-photon and *in vivo* microendoscopic Ca^2+^ imaging stacks are processed in slightly different ways to optimize the analysis of Ca^2+^ signals acquired in these different preparations (Fig. 2a; Supplementary Manual). The softwares described in this report are cross-platform Free and Open Source Software (FOSS) and licensed under the GNU GPLv3+ license (available at: *https://gitlab.com/d5674/occam*).

#### Preprocessing

*noise correction and ROI definition*. The input for Occam is an oligodendroglial Ca^2+^ imaging stack as recorded by the microscopy acquisition software. The Occam software initially performs noise correction steps according to the imaging condition, and then proceeds to an automatic ROI designation (Supplementary Manual). ROI designation is carried out using the machine learning-based WEKA Fiji/ImageJ2 plugin that performs a trainable segmentation of frames (Arganda-Carreras et al., 2017) combined with a local maxima segmentation tool. A ROI as designated by Occam is defined as a vector containing the mean fluorescence intensity value of the corresponding region in the different frames of the stack (that is, over the acquisition time points). Such ROI vectors are indifferently called ROI traces in this report. ROIs are sorted in two different classes: 1) high and medium mean pixel intensity ROIs, that correspond to Ca^2+^ active regions of the stack and 2) low mean pixel intensity ROIs, that are considered as background. In this report, WEKA classifiers were trained on Ca^2+^ imaging stacks obtained from GCaMP-expressing oligodendroglia in demyelinated lesions of the mouse corpus callosum in one-photon, two-photon and in vivo imaging conditions, separately. Notably, training WEKA is straightforward using any given set of image stacks. Occam’s user interface allows one to configure various aspects of the processing. For each processed image stack, Occam produces a set of comma-separated value (CSV) files that contain the description of the mean fluorescence intensity of each ROI along the time-resolved acquisition experiment. These files are then fed to the post-prOccam software.

#### Post-processing

*automated ROI refinement and quantifications*. The post-prOccam software is a configurable software that processes the files produced by Occam. Before performing a baseline subtraction, false positive ROI traces of a stack are rejected by using a sliding window-based subtraction of intensities in each ROI trace and applying a detection threshold defined by the mean absolute deviation (MAD) value on each subtracted trace (Supplementary Manual). Only ROIs that show significant Ca^2+^-fluorescence fluctuations are accepted for further analyses. A number of parameters configuring the ROI processing and the filtering stringency can be set in a configuration file (Supplementary Files 1-3). The accepted ROIs of a stack are output to a file and calculations that are then computed on these ROIs are output to another file (such as the surface area of each ROI in pixels, the ROI integral, the sum integral of all ROIs, the total ROI surface area in pixels, the sum integral corrected by the surface area of all ROIs, the Pearson correlation between each ROI and every other ROI; Supplementary Manual). Of note, this analysis can be performed on specific acquisition time point intervals of ROI traces by listing desired intervals in the configuration file. The post-prOccam software logs all the processing steps and their outcome to a file so as to let the user scrutinize the inner workings of the program.

### Statistical analysis

Data are expressed as mean ± SEM. GraphPad Prism (version 9.3.0; GraphPad Software Inc., USA) was used for statistical analysis. Each group of data was first subjected to Shapiro-Wilk normality test. According to the data structure, two-group comparisons were performed using the two-tailed unpaired Student’s t-test or the non-parametric two-tailed unpaired Mann-Whitney U test for independent samples; the two-tailed paired Student’s t-test was used for paired samples. Multiple comparisons were done with a two-way ANOVA test followed by a Bonferroni’s multiple comparison test.

### Data availability

The data generated and analyzed during this study are included in the manuscript and supplementary files. Occam and post-prOccam programs as well as all the software documentation and a full example of ex vivo widefield imaging data are hosted at *https://gitlab.com/d5674/occam* and published under a Free Software GNU GPLv3+ license.

## Supporting information

Supplementary Manual

## Acknowledgements

We thank the NeurImag platform and the animal facility of IPNP and their funding sources (Fédération pour la Recherche Médicale, Fondation Leducq). We would also like to thank Serge Charpak and Yannick Goulam for their help with the two-photon microscope, and Callum White for his contribution to initial data analysis in Python. This work was supported by grants from a subaward agreement from the University of Connecticut with funds provided by Grant No. RG-1612-26501 from National Multiple Sclerosis Society (NMSS), Fondation pour l’aide à la recherche sur la Sclérose en Plaques (ARSEP), Fondation pour la Recherche Médicale (FRM, EQU202103012626), ANR under the frame of the European Joint Programme on Rare Diseases (EJP RD, project no. ANR-19-RAR4-008-03) and ANR CoLD (ANR, ANR-20-CE16–0001-01). D.A.M. received a postdoctoral fellowship from Fondation pour la Recherche Medicale (FRM, project SPF202005011919) and a L’Oréal-UNESCO young talents award 2021 for women in science, B. M.-S. received a PhD fellowship from Université Paris Cité, C.H. received a postdoctoral fellowship from ARSEP. M.C.A. and F.R. are CNRS (Centre National de la Recherche Scientifique) investigators.

## Competing interests

The authors have declared that no competing interests exist.

## Author contributions

D.A.M. and B. M-S. conducted one-photon Ca^2+^ imaging experiments. C.H. performed two-photon Ca^2+^ imaging experiments and D.A.M. performed *in vivo* microendoscopy Ca^2+^ imaging experiments. D.A.M., B. M-S., C.H. and M.C.A. designed experiments and analysis. P.B., D.A.M., B. M-S wrote the ImageJ plugin. M.C.A. and F.R. designed the Python software and F.R. wrote the code. D.A.M. and M.C.A. performed data analyses and D.A.M., F.R. and M.C.A. wrote the manuscript. M.C.A supervised the project.

## Notes

https://gitlab.com/d5674/occam

### Competing Interest Statement

The authors have declared no competing interest.

## References

Abiraman, K., Pol, S. U., O’Bara, M. A., Chen, G.-D., Khaku, Z. M., Wang, J., Thorn, D., Vedia, B. H., Ekwegbalu, E. C., Li, J.-X., Salvi, R. J., & Sim, F. J. (2015). Anti-muscarinic adjunct therapy accelerates functional human oligodendrocyte repair. The Journal of Neuroscience: The Official Journal of the Society for Neuroscience, 35(8), 3676–3688. https://doi.org/10.1523/JNEUROSCI.3510-14.2015

Agarwal, A., Wu, P.-H., Hughes, E. G., Fukaya, M., Tischfield, M. A., Langseth, A. J., Wirtz, D., & Bergles, D. E. (2017). Transient Opening of the Mitochondrial Permeability Transition Pore Induces Microdomain Calcium Transients in Astrocyte Processes. Neuron, 93(3), 587-605.e7. https://doi.org/10.1016/j.neuron.2016.12.034

Bakiri, Y., Káradóttir, R., Cossell, L., Attwell, D. (2011). Morphological and electrical properties of oligodendrocytes in the white matter of the corpus callosum and cerebellum. J Physiol. 1;589(Pt 3):559–73. https://doi.org/10.1113/jphysiol.2010.201376.

Balia, M., Benamer, N., Angulo, M.C. (2017). A specific GABAergic synapse onto oligodendrocyte precursors does not regulate cortical oligodendrogenesis. Glia, 65(11):1821–1832. https://doi.org/10.1002/glia.23197.

Baraban, M., Koudelka, S., & Lyons, D. A. (2018). Ca (2+) activity signatures of myelin sheath formation and growth in vivo. Nature Neuroscience, 21(1), 19–23. PubMed. https://doi.org/10.1038/s41593-017-0040-x

Battefeld, A., Popovic, M. A., de Vries, S. I., & Kole, M. H. P. (2019). High-Frequency Microdomain Ca(2+) Transients and Waves during Early Myelin Internode Remodeling. Cell Reports, 26(1), 182-191.e5. PubMed. https://doi.org/10.1016/j.celrep.2018.12.039

Cai, D.J., Aharoni, D., Shuman, T., Shobe, J., Biane, J., Song, W., Wei, B., Veshkini, M., La-Vu, M., Lou, J., Flores, S.E., Kim, I., Sano, Y., Zhou, M., Baumgaertel, K., Lavi, A., Kamata, M., Tuszynski, M., Mayford, M., Golshani, P., Silva, A.J. (2016). A shared neural ensemble links distinct contextual memories encoded close in time. Nature, 2;534(7605):115–8. https://doi.org/10.1038/nature17955.

Cantu, D. A., Wang, B., Gongwer, M. W., He, C. X., Goel, A., Suresh, A., Kourdougli, N., Arroyo, E. D., Zeiger, W., & Portera-Cailliau, C. (2020). EZcalcium: Open-Source Toolbox for Analysis of Calcium Imaging Data. Frontiers in Neural Circuits, 14, 25. https://doi.org/10.3389/fncir.2020.00025

Cayre, M., Falque, M., Mercier, O., Magalon, K., Durbec, P. (2021). Myelin Repair: From Animal Models to Humans. Front Cell Neurosci, 14;15:604865. https://doi.org/10.3389/fncel.2021.604865.

Chittajallu, R., Aguirre, A., Gallo, V. (2004). NG2-positive cells in the mouse white and grey matter display distinct physiological properties. J Physiol. 15;561(Pt 1):109–22. https://doi.org/10.1113/jphysiol.2004.074252.

Chua, Y., & Morrison, A. (2016). Effects of Calcium Spikes in the Layer 5 Pyramidal Neuron on Coincidence Detection and Activity Propagation. Frontiers in Computational Neuroscience, 10. https://doi.org/10.3389/fncom.2016.00076

Cohen, R. I., & Almazan, G. (1994). Rat oligodendrocytes express muscarinic receptors coupled to phosphoinositide hydrolysis and adenylyl cyclase. The European Journal of Neuroscience, 6(7), 1213–1224. https://doi.org/10.1111/j.1460-9568.1994.tb00620.x

De Angelis, F., Bernardo, A., Magnaghi, V., Minghetti, L., & Tata, A. M. (2012). Muscarinic receptor subtypes as potential targets to modulate oligodendrocyte progenitor survival, proliferation, and differentiation. Developmental Neurobiology, 72(5), 713–728. https://doi.org/10.1002/dneu.20976

Deshmukh, V. A., Tardif, V., Lyssiotis, C. A., Green, C. C., Kerman, B., Kim, H. J., Padmanabhan, K., Swoboda, J. G., Ahmad, I., Kondo, T., Gage, F. H., Theofilopoulos, A. N., Lawson, B. R., Schultz, P.G., & Lairson, L. L. (2013). A regenerative approach to the treatment of multiple sclerosis. Nature, 502(7471), 327–332. https://doi.org/10.1038/nature12647

Duncan, I. D., & Radcliff, A. B. (2016). Inherited and acquired disorders of myelin: The underlying myelin pathology. Experimental Neurology, 283(Pt B), 452–475. https://doi.org/10.1016/j.expneurol.2016.04.002

Franklin, R. J. M., & Ffrench-Constant, C. (2017). Regenerating CNS myelin—From mechanisms to experimental medicines. Nature Reviews. Neuroscience, 18(12), 753–769. https://doi.org/10.1038/nrn.2017.136

Giovannucci, A., Friedrich, J., Gunn, P., Kalfon, J., Brown, B. L., Koay, S. A., Taxidis, J., Najafi, F., Gauthier, J. L., Zhou, P., Khakh, B. S., Tank, D. W., Chklovskii, D. B., & Pnevmatikakis, E. A. (2019). CaImAn an open source tool for scalable calcium imaging data analysis. ELife, 8, e38173. https://doi.org/10.7554/eLife.38173

Green, A. J., Gelfand, J. M., Cree, B. A., Bevan, C., Boscardin, W. J., Mei, F., Inman, J., Arnow, S., Devereux, M., Abounasr, A., Nobuta, H., Zhu, A., Friessen, M., Gerona, R., von Büdingen, H. C., Henry, R. G., Hauser, S. L., & Chan, J. R. (2017). Clemastine fumarate as a remyelinating therapy for multiple sclerosis (ReBUILD): A randomised, controlled, double-blind, crossover trial. Lancet (London, England), 390(10111), 2481–2489. https://doi.org/10.1016/S0140-6736(17)32346-2

Ingiosi, A.M., Hayworth, C.R., Harvey, D.O., Singletary, K.G., Rempe, M.J., Wisor, J.P., Frank, M.G. (2020). A Role for Astroglial Calcium in Mammalian Sleep and Sleep Regulation. Curr Biol, 16;30(22):4373-4383.e7. https://doi.org/10.1016/j.cub.2020.08.052.

Kirischuk, S., Scherer, J., Möller, T., Verkhratsky, A., & Kettenmann, H. (1995). Subcellular heterogeneity of voltage-gated Ca2+ channels in cells of the oligodendrocyte lineage. Glia, 13(1), 1–12. https://doi.org/10.1002/glia.440130102

Koizumi, S. (2010). Synchronization of Ca2+ oscillations: involvement of ATP release in astrocytes. FEBS J, 277(2):286–92. https://doi.org/10.1111/j.1742-4658.2009.07438.x.

Krasnow, A. M., Ford, M. C., Valdivia, L. E., Wilson, S. W., & Attwell, D. (2018). Regulation of developing myelin sheath elongation by oligodendrocyte calcium transients in vivo. Nature Neuroscience, 21(1), 24–28. PubMed. https://doi.org/10.1038/s41593-017-0031-y

Li, J., Miramontes, T., Czopka, T., Monk, K. (2022). Synapses in oligodendrocyte precursor cells are dynamic and contribute to Ca2+ activity. bioRxiv 2022.03.18.484955. https://doi.org/10.1101/2022.03.18.484955

Maas, D.A., Angulo, M.C. (2021). Can Enhancing Neuronal Activity Improve Myelin Repair in Multiple Sclerosis? Front Cell Neurosci, 15:645240. https://doi.org/10.3389/fncel.2021.645240.

Marisca, R., Hoche, T., Agirre, E., Hoodless, L. J., Barkey, W., Auer, F., Castelo-Branco, G., & Czopka, T. (2020). Functionally distinct subgroups of oligodendrocyte precursor cells integrate neural activity and execute myelin formation. Nature Neuroscience, 23(3), 363–374. https://doi.org/10.1038/s41593-019-0581-2

Moyon, S., Dubessy, A.L., Aigrot, M.S., Trotter, M., Huang, J.K., Dauphinot, L., Potier, M.C., Kerninon, C., Melik Parsadaniantz, S., Franklin, R.J., Lubetzki, C. (2015). Demyelination causes adult CNS progenitors to revert to an immature state and express immune cues that support their migration. J Neurosci, 7;35(1):4–20. https://doi.org/10.1523/JNEUROSCI.0849-14.2015.

Nualart-Marti, A., Solsona, C., & Fields, R. D. (2013). Gap junction communication in myelinating glia. Biochimica Et Biophysica Acta, 1828(1), 69–78. https://doi.org/10.1016/j.bbamem.2012.01.024

Paez, P. M., & Lyons, D. A. (2020). Calcium Signaling in the Oligodendrocyte Lineage: Regulators and Consequences. Annual Review of Neuroscience, 43, 163–186. https://doi.org/10.1146/annurev-neuro-100719-093305

Parys, B., Côté, A., Gallo, V., De Koninck, P., & Sík, A. (2010). Intercellular calcium signaling between astrocytes and oligodendrocytes via gap junctions in culture. Neuroscience, 167(4), 1032–1043. https://doi.org/10.1016/j.neuroscience.2010.03.004

Pitman, K. A., & Young, K. M. (2016). Activity-dependent calcium signalling in oligodendrocyte generation. The International Journal of Biochemistry & Cell Biology, 77(Pt A), 30–34. PubMed. https://doi.org/10.1016/j.biocel.2016.05.018

Rash, J. E., Yasumura, T., Dudek, F. E., & Nagy, J. I. (2001). Cell-specific expression of connexins and evidence of restricted gap junctional coupling between glial cells and between neurons. The Journal of Neuroscience: The Official Journal of the Society for Neuroscience, 21(6), 1983–2000.

Remaud, S., Ortiz, F.C., Perret-Jeanneret, M., Aigrot, M.S., Gothié, J.D., Fekete, C., Kvárta-Papp, Z., Gereben, B., Langui, D., Lubetzki, C., Angulo MC, Zalc B, Demeneix B. (2017). Transient hypothyroidism favors oligodendrocyte generation providing functional remyelination in the adult mouse brain. Elife, 6:e29996. https://doi.org/10.7554/eLife.29996.

Sahel, A., Ortiz, F. C., Kerninon, C., Maldonado, P. P., Angulo, M. C., & Nait-Oumesmar, B. (2015). Alteration of synaptic connectivity of oligodendrocyte precursor cells following demyelination. Frontiers in Cellular Neuroscience, 9, 77–77. https://doi.org/10.3389/fncel.2015.00077

Schindelin, J., Arganda-Carreras, I., Frise, E., Kaynig, V., Longair, M., Pietzsch, T., Preibisch, S., Rueden, C., Saalfeld, S., Schmid, B., Tinevez, J.-Y., White, D. J., Hartenstein, V., Eliceiri, K., Tomancak, P., & Cardona, A. (2012). Fiji: An open-source platform for biological-image analysis. Nature Methods, 9(7), 676–682. https://doi.org/10.1038/nmeth.2019

Shuman, T., Aharoni, D., Cai, D.J., Lee, C.R., Chavlis, S., Page-Harley, L., Vetere, L.M., Feng, Y., Yang, C.Y., Mollinedo-Gajate, I., Chen, L., Pennington, Z.T., Taxidis, J., Flores, S.E., Cheng, K., Javaherian, M., Kaba, C.C., Rao, N., La-Vu, M., Pandi, I., Shtrahman, M., Bakhurin, K.I., Masmanidis, S.C., Khakh, B.S., Poirazi, P., Silva, A.J., Golshani, P. (2020). Breakdown of spatial coding and interneuron synchronization in epileptic mice. Nat Neurosci, 23(2):229-238. https://doi.org/10.1038/s41593-019-0559-0.

Takeda, M., Nelson, D. J., & Soliven, B. (1995). Calcium signaling in cultured rat oligodendrocytes. Glia, 14(3), 225–236. https://doi.org/10.1002/glia.440140308

Venugopal, S., Srinivasan, R., & Khakh, B. S. (2019). GECIquant: Semi-automated Detection and Quantification of Astrocyte Intracellular Ca2+ Signals Monitored with GCaMP6f. In M. De Pittà & H. Berry (Eds.), Computational Glioscience (pp. 455–470). Springer International Publishing. https://doi.org/10.1007/978-3-030-00817-8_17

Wang, Y., DelRosso, N. V., Vaidyanathan, T. V., Cahill, M. K., Reitman, M. E., Pittolo, S., Mi, X., Yu, G., & Poskanzer, K. E. (2019). Accurate quantification of astrocyte and neurotransmitter fluorescence dynamics for single-cell and population-level physiology. Nature Neuroscience, 22(11), 1936– 1944. https://doi.org/10.1038/s41593-019-0492-2

Wang, Y., Shi, G., Miller, D. J., Wang, Y., Wang, C., Broussard, G., Wang, Y., Tian, L., & Yu, G. (2017). Automated Functional Analysis of Astrocytes from Chronic Time-Lapse Calcium Imaging Data. Frontiers in Neuroinformatics, 11, 48. https://doi.org/10.3389/fninf.2017.00048

Welliver, R. R., Polanco, J. J., Seidman, R. A., Sinha, A. K., O’Bara, M. A., Khaku, Z. M., Santiago González, D. A., Nishiyama, A., Wess, J., Feltri, M. L., Paez, P. M., & Sim, F. J. (2018). Muscarinic Receptor M _3_ R Signaling Prevents Efficient Remyelination by Human and Mouse Oligodendrocyte Progenitor Cells. The Journal of Neuroscience, 38(31), 6921–6932. https://doi.org/10.1523/JNEUROSCI.1862-17.2018

Xu, Y. K. T., Call, C. L., Sulam, J., & Bergles, D. E. (2021). Automated in vivo Tracking of Cortical Oligodendrocytes. Frontiers in Cellular Neuroscience, 15, 667595. https://doi.org/10.3389/fncel.2021.667595

Yuan merge;2015 Tianyi Yuan, T., Lu, J., Zhang, J., Zhang, Y., Chen, L. (2015). Spatiotemporal detection and analysis of exocytosis reveal fusion “hotspots” organized by the cytoskeleton in endocrine cells. Biophysical journal, 108(2), 251–260. https://doi.org/10.1016/j.bpj.2014.11.3462

Zhang, L., Liang, B., Barbera, G., Hawes, S., Zhang, Y., Stump, K., Baum, I., Yang, Y., Li, Y., Lin, D.T. (2019). Miniscope GRIN Lens System for Calcium Imaging of Neuronal Activity from Deep Brain Structures in Behaving Animals. Curr Protoc Neurosci, 86(1):e56. https://doi.org/10.1002/cpns.56.

