## Supplementary Manual for "Versatile and automated workflow for the analysis of oligodendroglial calcium signals in preclinical mouse models of myelin repair"

### Manual of Occam & post-prOccam

*Release 1.0 (20221014)*

**Oct 14, 2022**

CHAPTER

ONE

### LEGAL NOTICE

Copyright 2022, Filippo Rusconi, Philippe Bun, Dorien Maas, Maria Cecilia Angulo

This user manual is licensed under the GNU GPLv3+ license, exactly as is the whole software package it documents. See the LICENSE file in the distribution.

CHAPTER

TWO

### GENERALITIES

#### 2.1 Introduction

This documentation project is in support of an experimental work performed in Maria Cecilia Angulo's team laboratory in IPNP, Paris.

There are two user manuals in this project, each documenting one of the two software projects that have been developed in the context of that experimental work: the manuals for the `Occam` and the `post-prOccam` programs are documented in the order that the programs are put at work in the experimental work.

The directory named `example-data` in the documentation source tree contains one zip-based archive

- `ex-vivo-widefield.7z`: data from an ex vivo widefield project;

**Note:** The figures in the `Occam` user manual are taken from the `ex-vivo-widefield` project.

These archives not only contain all the files that are needed to run both `Occam` and `post-prOccam` but also contain all the result files that were produced upon running both programs.

**Warning:** The data in the example directory are large (in the Gb size range) and are added to the Git repository under the Git LFS framework. They are not meant to be modified and committed to the Git repository in any way.

The `post-prOccam` software runs with a configuration file. The `configuration-examples` directory contains three sample configuration files found to work best in the three experimental setups:

- `ex-vivo-widefield.cfg`
- `ex-vivo-2-photon.cfg`
- `in-vivo-widefield.cfg`

#### 2.2 Installation

As portable software, `Occam` and `post-prOccam` can be used on GNU/Linux and MS Windows platforms. The installation procedure, along with the system requirements are described for these two platforms.

##### 2.2.1 System pre-requisites

- The `Occam` software is a `Fiji/ImageJ2` plugin; hence it requires this software, that can be downloaded from the official repository. Please, take note of the installation directory, as the `plugins` directory inside it will be needed later.

  The `Weka` segmentation plugin for `Fiji/ImageJ2` is required and can be downloaded from this site.

- The `post-prOccam` software was developed on Debian GNU/Linux (either stable or testing). The main pre-requisite for running `post-prOccam` is to have Python3 installed (lowest functional version is 3.9.2).

  GNU/Linux

  On a desktop computer where Python3 is already installed, the extra Python3 requirements are easily fulfilled using the following command (as root):

  ```
  # apt install python3-pandas python3-regex python3-matplotlib
  ```

  MS Windows

  Python3 might be installed in two ways:

  - By installing the Anaconda environment, which is huge and might very well be overkill for our `post-prOccam` software program. If the user has the environment already installed, then nothing more should be needed.
  - By installing Python from scratch, according to the following steps:
    * Install Python using an installer such as `python-3.10.8-amd64.exe` (this is the one we used to test the software on a clean machine prior to making the first public release). The installer was downloaded from this official site.

**Note:** In our setup, we asked that the Python.exe interpreter be added to the PATH for *all* users. This ensures that typing `python.exe` in a console finds the interpreter just fine.

Also, the package was installed at `C:\Program files\Python310`.

To check this, in a `cmd.exe` Windows terminal, issue the following command:

echo %PATH%

```
C:\Program Files\Python310\Scripts\;C:\Program Files\Python310\;
[skip other paths]
```

The PATH indeed includes the relevant Python-related paths.

* The following `pip`-based commands need to be issued in order to install the corresponding requirements:

```
> pip install pandas
> pip install matplotlib
> pip install regex
```

#### 2.3 Download and install the software

The software is licensed under the GNU GPLv3+ Free Software license (see the `LICENSE` file in the distribution) and is available at the GitLab repository.

There are two different ways to install the project:

- As a `zip` archive by browsing to this GitLab page. Extracting the archive creates the `occam-main` directory.
- As a Git repository by cloning the repository:

  `git clone https://gitlab.com/d5674/occam.git`

  or

  `git clone git@gitlab.com:d5674/occam.git`

> **Warning:** The cloning will take some time to complete even after the `Updating files: 100% (253/253), done.` line is printed because there is a big example-data archive in the repository. Please, do not interrupt the cloning operation until it is actually finished, that is, when the git program gives back the prompt.

#### 2.4 Code, documentation and example data

The project's main directory contains two main subdirectories:

- `devel`: the directory that contains the code (`Occam` and `post-prOccam` subdirectories);
- `doc`: the documentation directory (containing the `example-data` subdirectory).

The `devel/Occam/Occam_.ijm` Fiji/ImageJ2-based plugin is to be copied to the `plugins` directory in the main installation directory of `Fiji/ImageJ2`. Once this is done, run the `Fiji/ImageJ2` program and follow the instructions in the `Occam` user manual in the next chapters. Note that the example data discussed in the following paragraphs are useful to test the setup.

The `devel/post-prOccam/post-prOccam` file is the main Python file to be executed (that is, interpreted using the `Python.exe` (MS Windows) or `python3` (GNU/Linux)) interpreter).

The `doc` directory contains example data that can be used to test the setup. After having cloned the Git repository (or extracted the archive), that `doc` directory contains the `example-data` subdirectory which contains the `ex-vivo-widefield.7z` archive. Please, use 7zip to extract it in place so that the obtained directory is located at `doc/example-data/ex-vivo-widefield`.

That directory now contains three elements:

- The `ex-vivo-widefield-stack.tif` frame stack to use to test `Occam`;
- The `Weka` classifier model `classifier-calcium-ex-vivo-widefield.model`;
- The `ex-vivo-widefield-stack_results` directory containing the files generated by `Occam` (do not overwrite them, make a pristine copy of this directory before running `Occam`) to test `post-prOccam`.

At this point testing the setup involves running the `Occam` program on the `ex-vivo-widefield-stack.tif` image stack, as described in the user manual for that program.

Next, the `post-prOccam` software might be tested by running it from inside the `ex-vivo-widefield-stack_results` directory, as described in the user manual for that program.

CHAPTER

THREE

### OCCAM MANUAL

#### 3.1 Introduction

In this chapter, we describe the `Fiji/ImageJ2`-based `Occam` plugin (Oligodendroglial cells calcium activity monitoring) that we developed to analyze $Ca^{2+}$ signals of GCaMP-expressing oligodendroglia in demyelinated lesions. `Occam` runs on wide-field ex vivo, 2-photon ex vivo and in vivo microendoscopy $Ca^{2+}$ imaging stacks. This program has the following runtime requirements:

- For this release, `Occam` was tested with `Fiji/ImageJ2` version 1.53t and also worked with earlier versions;
- The `Weka` `Fiji/ImageJ2` plugin with a user-trained `Weka` classifier. A `Weka` classifier model for the ex vivo widefield microscopy experimental setup is provided as file `classifier-calcium-ex-vivo-widefield.model` in the `devel/Occam` directory of the distribution.

#### 3.2 Running Occam¶

To run `Occam`, save the plugin file in the `Fiji/ImageJ2` plugin folder, start `Fiji/ImageJ2` and launch the `Occam` plugin.

**Note:** Note that the `Occam` plugin file is named `Occam_.ijm`. That name should not be changed because otherwise the plugin will now show up in the *Plugins* menu in `Fiji/ImageJ2`.

Once launched, a dialog box with four user entries needs to be filled in:

- The type of experiment. Here the user has to choose one of out three options: ex vivo (widefield), ex vivo (2-photon) and in vivo (widefield);
- The `Weka` classifier file location;
- The minimum size for an active region to be considered (square pixels);
- Only applicable for in vivo imaging: the minimum percentage of pixel overlap for detected ROIs in different substacks in the same region of the frame.

By default, the minimum size of an active region has an area of 300 pixels (that is, if that region were square, it would have ca 17-pixel-long sides) which corresponds to the value used in our analysis of widefield ex vivo $Ca^{2+}$ imaging data. The minimum percentage of overlapped ROI area that allows the merging of ROIs from different substacks is automatically set to 15%, which corresponds to the value used in our analysis of in vivo data. However, both these values can be adjusted by the user.

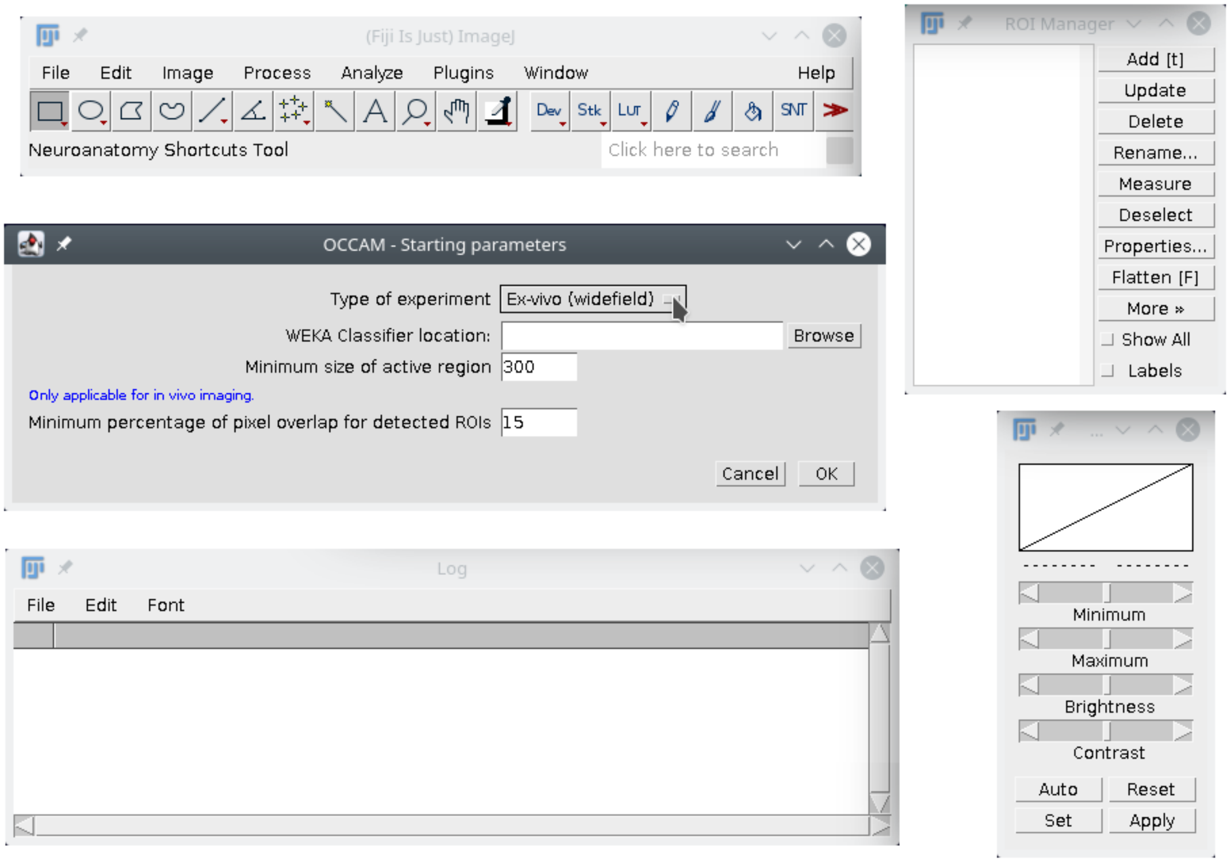

Fig. 3.1: `Occam` - Starting parameters

##### 3.2.1 Step-by-step procedure¶

The `Occam` plugin performs different operations in six different steps and creates a number of files.

###### Step 1: Preparation of the time-resolved frame stack

Each fluorescent time series (that is, a stack of time-resolved frames) is first converted to a series of 8-bit images. The images are further binned by a factor 2 if they are wider than 1920 pixels. At this point, a dialog window titled "Space and time parameters for the stack processing" opens to offer three options, as shown in Fig. 3.2 below.

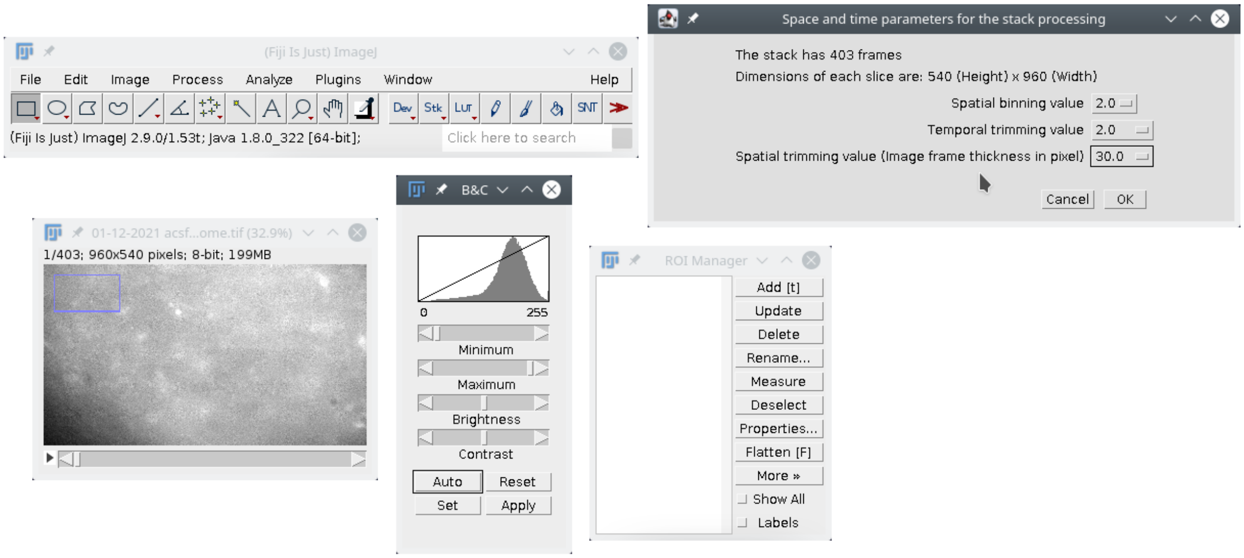

Fig. 3.2: `Occam` - Space and time parameters for the processing

- The spatial binning value allows for an additional binning of images in the stack if desired;
- The temporal trimming value allows to remove the same number of frames at the beginning and at the end of the stack;
- The spatial trimming value allows all the frames to be cropped by removing an outer frame having the width of the designated number of pixels. This procedure can be useful in the case of potential edge illumination artifacts.

For ex vivo imaging data, the resulting frame stack is saved in a file named `no-bleach-time-trimmed-space-trimmed-binned.tif` for further processing (Fig. 3.3). For in vivo imaging data, an additional dialog box makes it possible to reduce the number of images per stack, when this is too large, by averaging every pair of frames. The resulting frame stack is saved in the file `avgbin-no-bleach-no-trim-no-bin-downsampled.tif` for further processing.

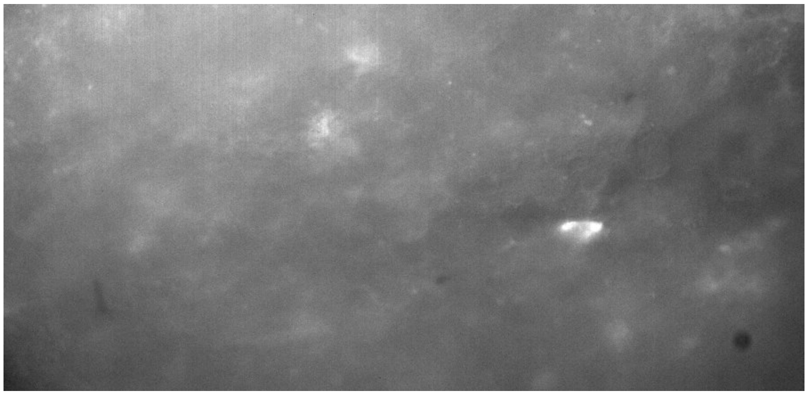

Fig. 3.3: Occam - Example of a no-bleach-time-trimmed-space-trimmed-binned.tif image obtained from an ex vivo widefield stack

##### Step 2: Photobleaching correction¶

The photobleaching correction applies only on stacks of ex vivo (widefield) experiments because they exhibit a greater photobleaching than that observed in other experiments. This option is therefore skipped for 2-photon ex vivo and widefield in vivo experiments.

To correct for photobleaching, Occam derives the correction parameters by fitting the mean fluorescence intensity over time with a double exponential decay curve:

y = A + B*exp(-C*t) + D*exp(-E*t)

where:

- A: the offset of the intensity;
- B, D: the amplitude of each exponential;
- C, E: the characteristic time decay of the t exponential.

The following constraints were applied: A > 1 and D > 0.

The ratio between the raw and the fitted mean fluorescence intensity is calculated to obtain the corrected mean fluorescence intensity over time. The procedure is repeated 33 times, each time removing a block of 34 images from a different part of the time series. At the end of the process, the user is presented with a series of graphs displaying the mean fluorescence intensity over time (red), the fit (blue), the corrected mean fluorescence intensity of the image stack where the fit applies (black) and the goodness of fit $R^2$ . The plots are saved as a stack in the fitting-profiles.tif file (see Fig. 3.4 below). The user can review this file and select the best fit that will be used for bleaching correction. After the photobleaching correction step, the fluorescence time-resolved frame stack is saved in the pb-corrected-stack.tif file. Note that the chosen fit will be labeled *Selected* in the fitting-profiles.tif file. In case not a single fit produces a reliable photobleaching correction, the user can choose to not perform any bleaching correction. In this case, the fluorescence time series is saved in the nopb-corrected-stack.tif file.

The customized photobleaching correction procedure described here was applied to our data because fitting a line on the complete z-axis profile often produces unreliable photobleaching corrections, because oligodendroglial cells $Ca^{2+}$ signals are characterized by very slow increase and decay times lasting several seconds (sometimes minutes) and therefore occupying a large part of the z-axis profile. Removing a part of the z-axis profile that includes such large $Ca^{2+}$ signals from the calculation of the fitted line produces a reliable photobleaching correction.

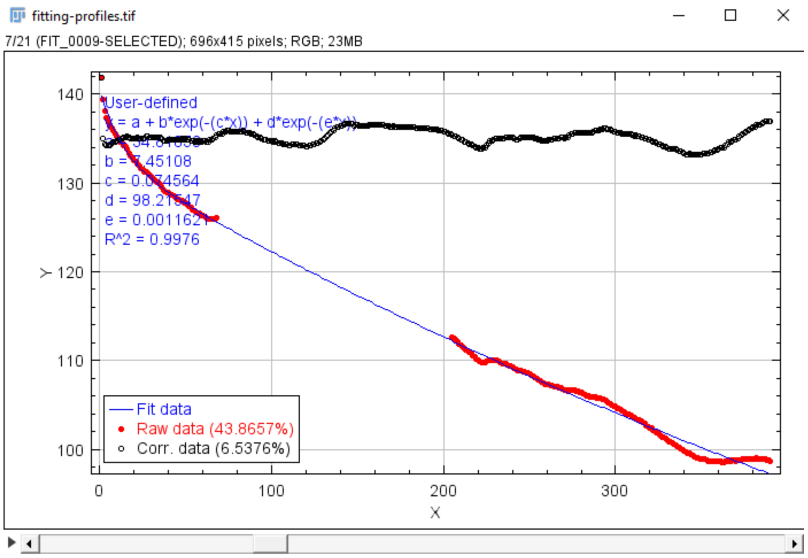

Fig. 3.4: `Occam` - Selected fit in the `fitting-profiles.tif` file obtained from the ex vivo widefield stack of Figure 1.

##### Step 3: Noise removal and projection for Weka¶

*For ex vivo imaging data (widefield and 2-photon)*

Removal of noise from the frame stack is performed using a combination of filters:

- A Fourier transform filter for the removal of frequencies within a 2-pixel-radius circle centered on (0,0);
- A rolling ball subtraction background filter of 500 pixels;
- A Gaussian blur with a sigma of 5 pixels.

Following noise removal, the sum of the maximum and the sum z-projections is generated and subjected to a median filter (radius 4.0). The resulting image is saved as `projection-for-WEKA.tif` and used as the input for the `Weka` classifier (Fig. 3.5).

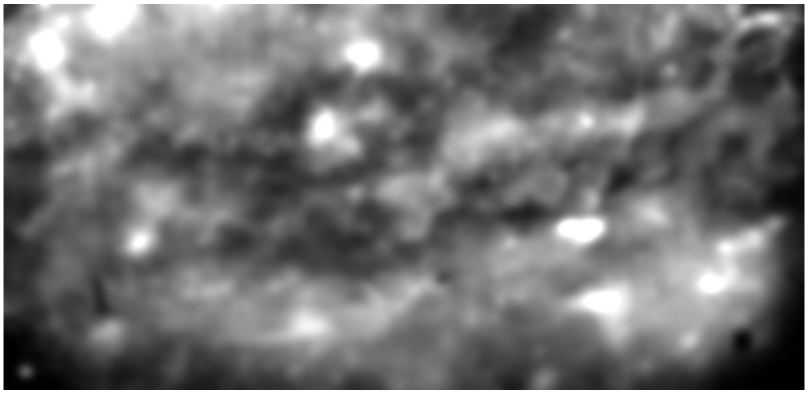

Fig. 3.5: `Occam` - Image saved in file `projection-for-WEKA.tif` for the example in Figure 1.

*For in vivo imaging data*

Because in vivo imaging typically involves the acquisition of image stacks over long periods of time —in our case 7 minutes (4200 frames)— the stacks need to be split into a number of frame groups (that is, sub-stacks). This grouping method ensures a more accurate detection of ROIs that are active only during short periods of time. To help the user determine a proper size of the sub-stacks (that is, the number of frames in each sub-stack), a list of possibilities is provided. The list is filled-in with sub-stack size values calculated by `Occam` to provide an integral number of sub-stacks.

**Note:** The user should be aware that a previous binning or trimming procedure may change the list of possible "number of frames". Once this number is selected, `Occam` splits the image stack into sub-stacks by dividing the total number of images by the selected number, and proceeds with the noise correction and projection for `Weka` as described above on each individual sub-stack.

Removal of noise from the sub-stacks is done by subtracting the minimum z-projection from each frame in the sub-stack followed by a rolling ball background subtraction with a filter of 20 pixels. Of note, no Fourier transform calculation is used in this case.

Following noise removal, the multiplication of the maximum and the standard deviation z-projections is subjected to a median filter (radius 4.0) and used as input for the `Weka` classifier.

##### Step 4: Weka segmentation¶

The `Weka` plugin (Arganda-Carreras et al., *Bioinformatics*, 2017) is a trainable segmentation software piece that we use to define four different types of ROIs:

- mean high pixel intensity;
- medium pixel intensity;
- mean low pixel intensity;
- mean background (considered as background signal).

The mean and medium pixel intensity ROIs are considered active ROIs from the $Ca^{2+}$ signals stand point, while mean low pixel intensity are not.

The user must train their own `Weka` classifier and load it in the user interface of the `Occam` `Fiji/ImageJ2` plugin.

**Note:** It is essential that the training be performed with images similar to that shown as an example in `projection-for-WEKA.tif`.

In our case, we trained three different `Weka` classifiers using $Ca^{2+}$ imaging frame stacks obtained in the three different types of experiments. During the `Weka` training, we used a fast random forest decision-making algorithm with the following features: Gaussian blur, Hessian, sober filter, membrane projections (thickness 1, patch size 5).

To get a first ROI segmentation, `Occam` loads both the `projection-for-WEKA.tif` file and the `Weka` classifier. Then, it runs the `Weka` segmentation of the image. The segmentation result is saved in the `results_weka.tif` file as an image showing each type of ROI in one color (Fig. 3.6; in our case, red and green designate high and medium pixel intensity ROIs and purple and yellow designate low pixel intensity and background ROIs). Note that for in vivo image analysis a `Weka` segmentation is run separately for each sub-stack with its corresponding `projection-for-WEKA.tif` file. In this case, the `results_weka.tif` file contains the same number of frames as the number of sub-stacks.

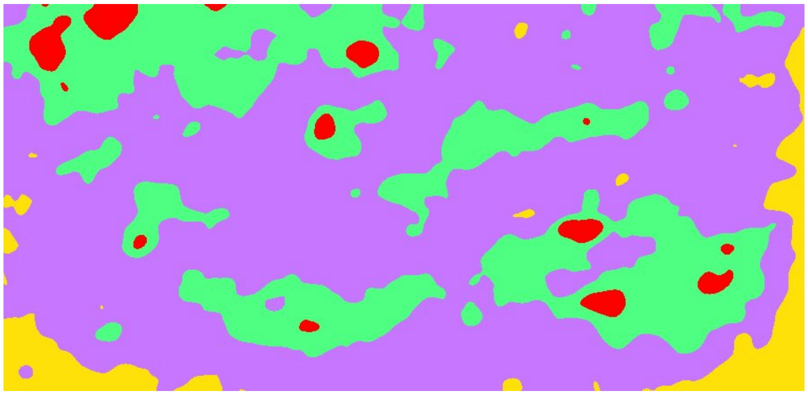

Fig. 3.6: `Occam` - Image of the `Weka` segmentation saved as `results_weka.tif` from the example in Figure 3.

##### Step 5: ROI refinement

To refine the `Weka`-based ROI designation, `Occam` uses the `projection-for-WEKA.tif` file and applies a local maxima algorithm and segmentation tool to identify areas that contain a local maximum. When multiple local maximum areas are located inside one `Weka`-determined ROI, they are considered as individual ROIs instead of the bigger `Weka`-generated ROI (Fig. 3.7). ROIs with an area smaller than a configurable number of pixels are discarded (the value is configurable in the dialog window when starting Occam).

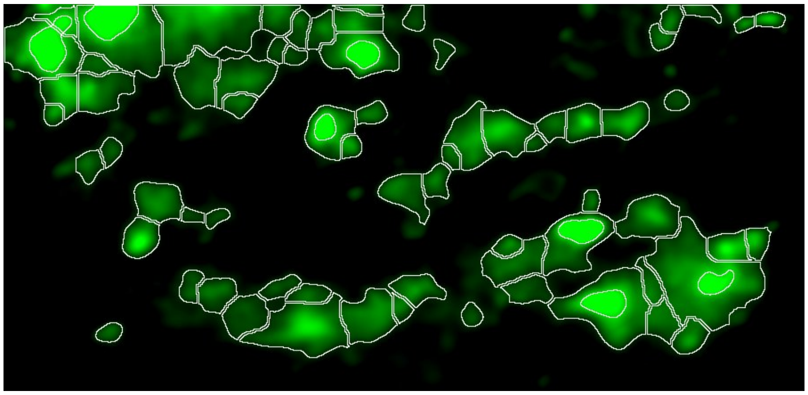

Fig. 3.7: `Occam` - All ROIs (white) detected in the stack of Fig. 3.1

*For in vivo imaging data*

Since the ROI designation and refinement is done separately for each sub-stack, each stack has a number of ROI sets that can attain the number of sub-stacks. It is important to note that a specific region of a stack could be active at different time points during an acquisition and, therefore, that this same region can be detected as a ROI in multiple sub-stacks, representing more than one ROI in the full stack. To overcome this problem, `Occam` projects all ROIs from all sub-stacks (contained in files `roi-high.zip` and `roi-medium.zip`) onto each other to determine the degree of overlap among all detected ROIs in a stack. The files `overlap_roi-medium.tif` and `overlap_roi-high.tif` contain a visualization of ROI overlaps using a pseudo color scale. To define if two or more overlapping ROIs actually correspond to a single ROI, the user configure a percentage of overlap beyond which the overlapping ROIs are considered as a single ROI (this value can be configured in the dialog window when starting Occam). One overlapping ROI is merged with its other overlapping ROIs if its percentage of overlap with the total area occupied by these ROIs is at least equal to the set value. The final ROI set is saved in the files `roi-high_final.zip` and `roi-medium_final.zip`.

##### Step 6. Data collection

Finally, several files containing the information of each ROI are generated:

- The `roi-high.zip` and `roi-medium.zip` archive files for ex vivo imaging data and the `roi-high_final.zip` and `roi-medium_final.zip` archive files for in vivo imaging data contain the ROI contours;
- The `roi-medium-mean-intensity.csv` and `roi-high-mean-intensity.csv` files containing mean pixel intensities of each detected ROI in a stack;
- The `roi-high-area-xy.csv` and `roi-medium-area-xy.csv` files containing the size in pixels of all the detected ROIs; These output files are used as input for the post-prOccam software.

Followed by the *post-prOccam* user manual.

CHAPTER

FOUR

### POST-PROCCAM MANUAL

#### 4.1 Introduction

In the previous *Occam* chapter, we described the `Fiji/ImageJ2`-based software that we developed to perform an unbiased and automated segmentation of active regions on $Ca^{2+}$ imaging stacks.

For each image stack, `Occam` does produce the following files that are located in the same directory that contained the image stack file (the data directory):

- `roi-high-mean-intensity.csv`
- `roi-high-area-xy.csv`
- `roi-medium-mean-intensity.csv`
- `roi-medium-area-xy.csv`

These four files constitute the base material on which `post-prOccam` operates.

**Note:** It might happen that `Occam` does not find any significant ROI in any of the frames of the image stack. In this case, some of the files above might not be produced. `post-prOccam` does check for this situation.

The files above are located inside a data directory that might have any name. For example, one of the data directories in our project has the following name:

`29-10-2021-m1s1r2-PDGFGcp6-nodrug-results`

The four files above are used by `post-prOccam` to perform data processing of both the following:

- the mean intensities of the high- and medium-intensity detected ROIs
- the mean areas of the high- and medium-intensity detected ROIs

The data processing that is carried over by `post-prOccam` is configured in a configuration file of which the path name needs to be provided on the command line. Sample configuration files suitable for different types of experiments are shipped along with this documentation in the `example-data/configuration-examples` directory. In the rest of this text the configuration file will be designated like so: `config.cfg`. The user is advised to use these files as starting points for the elaboration of new configurations that suit best their needs.

#### 4.2 Configuration of post-prOccam

The `config.cfg` file contains three main sections:

- *[MAIN]*
- *[INTEGRATION]*
- *[CORRELATIONS]*

---

**Note:** Any line in the configuration file that starts with '#' is a comment and is not interpreted by post-prOccam.

---

The configuration parameters in each section are detailed below.

##### 4.2.1 The [MAIN] section

- ACQ_RATE = 0.5716

  The time lag in seconds between two frames of the image stack

- FRAME_TOTAL_AREA = 432000

  The total area of the frame in square pixels, that is, the product of num_pix_horizontal * num_pix_vertical

- ROI_TRIM_INTERVALS =

  1-14
  # 348-362
  14 - LAST

  The ROI mean intensity series can be trimmed, typically at the front and at the back of the series. To define trimming intervals uncomment the lines and set appropriate values. The "14 - LAST" format is a convenience format to trim the last 14 rows of the ROI. Change the 14 value to any appropriate value. In the example above, the `# 348-362` line is not interpreted, as it is commented out using the # character.

  ---

  **Note:** Please, do not comment out the "ROI_TRIM_INTERVALS =" text line, because the program will fail if this line is commented out or absent from this configuration file. Only comment out the subsequent lines, because the value of that parameter can be empty, like so:

  ROI_TRIM_INTERVALS =

  # 1-14
  # 348-362
  # 14 - LAST

  ---

- MIN_MEAN_INTENSITY_HALF_WINDOW = 3

  The number of ROI points to be used to compute the minimum mean intensity of the ROI. If the parameter is set to 3, the minimum mean intensity is computed for the following window:

```
[ . . . x . . . ]
        |
        v
        lowest intensity value
```

3 points are selected on the left the of minimum intensity value `x` of the ROI and the same number of points are selected on the right of that minimum intensity value. The total window thus comprises 7 points, which are used to compute the minimum mean intensity value.

In the example above, if the minimum intensity point is found near the front (or the back) of the ROI series, the first 7 points (or last 7 ones) are used to compute the minimum mean intensity value.

- MEAN_ABS_DEVIATION_WINDOW = 40

  The width of the "sliding window" that is used to compute the ROI[z] - ROI[x] subtraction over all the ROI intensity values. In the example above, the ROI[z] - ROI[x] window comprises 40 points. In our case, this value is used to do a sliding window subtraction of points distant 40 frames which roughly match the mean rise time of oligodendroglia $Ca^{2+}$ events.

- MEAN_ABS_DEVIATION_FACTOR = 1.5

  Number by which the mean absolute deviation values are multiplied to yield the threshold governing the acceptance of the ROI in the final set of ROIs.

- CONDITION_MATCHING_MIN_POINT_COUNT = 6

  Number of points in each ROI that must have a mean value greater than the threshold (see previous parameter) in order for the ROI to be accepted.

##### 4.2.2 The [INTEGRATION] section

- INTERVALS =

#15-30 "no stim"
#70-120 "stim "
#200-300 "wash"

The ROI mean intensity integration can be performed on the whole ROI series or only on parts of that series. The values above —*if uncommented*— will elicit a range-based integration for the ROI in three "sections" of the ROI. The first section (*15-30*) elicits the integration over the corresponding ROI point range (each number describes the row in the ROI series, that is, the number of the frame in the image stack). The integration will be labelled in the processing report file using the text that follows the frame number range (note that the label text has to be in double quotes).

If the integration must be performed over the whole range of the ROI, then comment out (as shown above) all the intervals.

**Note:** Please, do not comment out the "INTERVALS =" text line, because the program will fail if this line is commented out or absent from this configuration file. Only comment out the subsequent lines, because the value of that parameter can be empty.

Also, do not comment out the "[INTEGRATION]" section line because the program will fail if this line is commented out or absent from this configuration file.

##### 4.2.3 The [CORRELATIONS] section

- PERFORM_ROI_CORRELATIONS = True

  Set to True if the inter-ROI Pearson correlations must be performed, otherwise set to False.

- ROI_CORRELATION_THRESHOLD = 0.9

  Threshold above which two ROIs are considered correlated; it corresponds to the Pearson correlation coefficient. This is an absolute value. ROIs will be considered correlated if their correlation value is either less than -ROI_CORRELATION_THRESHOLD (negative correlation) or greater than ROI_CORRELATION_THRESHOLD (positive correlation).

#### 4.3 Running post-prOccam

To run the `post-prOccam` program, enter the directory that contains the four files described above (that is, the data directory) and enter the following command line, with as single parameter the full path to the configuration file:

```
post-prOccam </path/to/the>/config.cfg
```

To have a help message describing the parameters to add to the command, enter

```
post-prOccam -h
```

---

**Note:** Make sure the `post-prOccam` program is in the PATH. Otherwise, call the program with its full path name:

- On MS Windows, provide the full pathname like so (adapt the path): `c:\users\<logname>\devel\oligo\post-prOccam`
- On GNU/Linux, provide the full pathname like so (adapt the path): `/home/<logname>/devel/oligo/post-prOccam`

---

`post-prOccam` will check the existence of the four files that are output by Occam. First, it checks the existence of the `high` input data file pair (`roi-high-mean-intensity.csv` and `roi-high-area-xy.csv` ), like so: if `roi-high-mean-intensity.csv` is found, then `roi-high-area-xy.csv` has to be found also (otherwise an exception is raised with a meaningful message to the user). If `roi-high-mean-intensity.csv` is `not` found, then the `high` data set is discarded altogether. The same identical logic is applied to the `medium` input data file pair (`roi-medium-mean-intensity.csv` and `roi-medium-area-xy.csv`).

#### 4.4 The files created by post-prOccam

During its execution, the `post-prOccam` program creates a number of files. These files are described in the following sections.

##### 4.4.1 The `log` file

The program logs all the debug messages issued during its execution in this file. The `log` file is also extremely useful to understand the inner workings of the program. It is packed with information that explains the state of the data while it is being processed.

##### 4.4.2 The `rois.csv` file

This file contains the set of *accepted* ROIs. Each ROI is represented as a column, with the ROI's name as the header of the column.

##### 4.4.3 The `stats.csv` file

This file contains all the results of the ROI analysis by `post-prOccam`.

##### 4.4.4 The `rois-plots.png` file

This file, along with its SVG and PDF format counterparts, contains the plots of the ROIs involved in the analysis. The plots are divided into two categories: High- and Medium-intensity plots. Also, the plots show all the accepted ROIs (upper panels) and the rejected ROIs (lower panel).

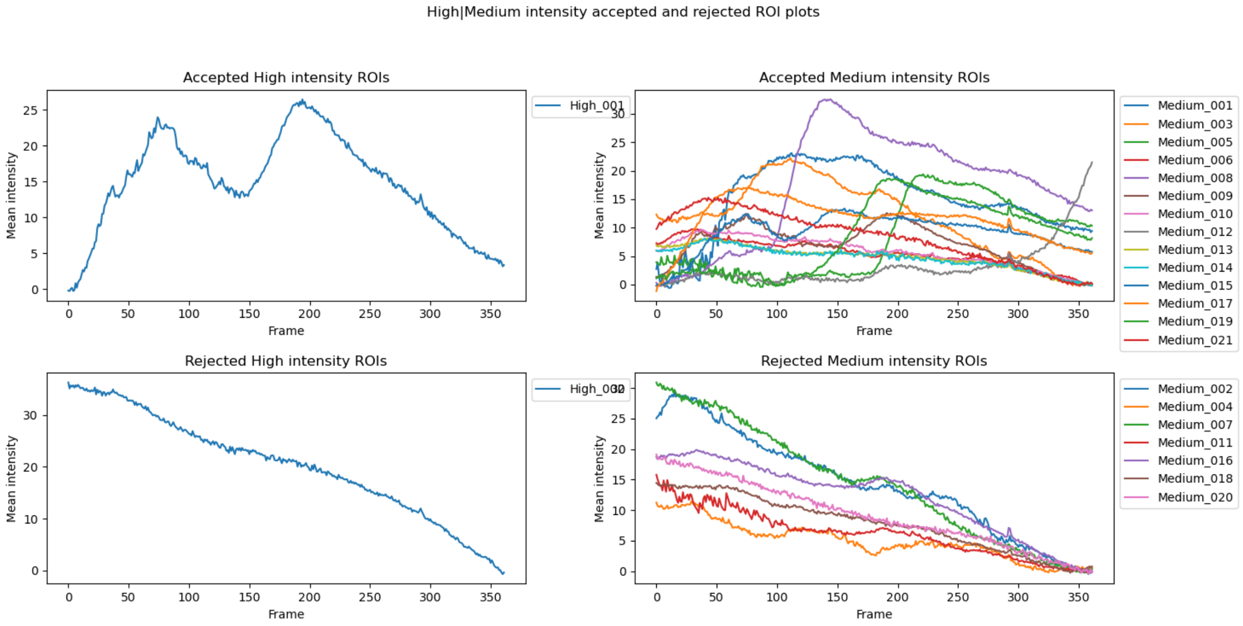

Fig. 4.1: Plots of the ROIs in the analyzed data set

##### 4.4.5 The `correlations-plots.png` file

This file, along with its SVG and PDF format counterparts, contains the plot of the inter-ROI Pearson correlations matrix performed only on the accepted ROIs.

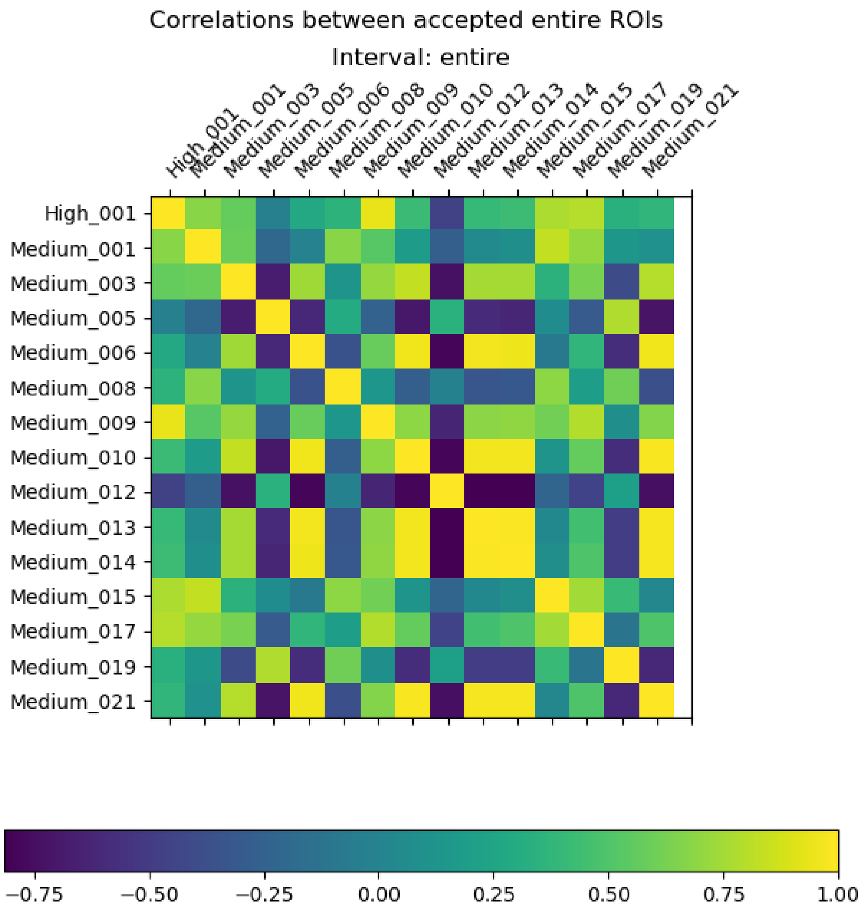

Fig. 4.2: Plot of the inter-ROI correlation matrix

#### 4.5 Detailed description of the program's operations

In this section, the various steps involved in the program's operations will be detailed so as to provide the user with a clear understanding of its inner workings.

##### 4.5.1 Parsing of the command line parameters

The program takes a number of parameters on the command line, as described *here*.

If the number of arguments is not as expected by the program, a help message is printed to the console.

##### 4.5.2 Loading of the `config.cfg` file.

The program reads the configuration file of which the path has to be provided on the command line as detailed in *the introduction*.

##### 4.5.3 Loading of the data in Pandas objects

The program uses the Pandas Python library to handle most of the data in this project and to perform a vast part of all the calculations.

The whole set of ROIs, loaded from the High and Medium mean intensity data files (`roi-high-mean-intensity.csv` and `roi-medium-mean-intensity.csv`), are set in a Pandas `data frame`. Each ROI in the data frame is like a column of values and is called a `data series`. There are as many data series in the data frame as there are ROI columns in the loaded files.

##### 4.5.4 Processing of the data

The differents steps of the data processing are detailed below.

###### Trimming of the front and back of the ROIs

The ROIs are trimmed to remove from all the ROI series the rows specified in the `config.cfg` file, in the variable `ROI_TRIM_INTERVALS`. Each row in each trim interval range specified in the configuration file is eliminated from the ROI series.

###### Filtering of the ROI data

We want to filter the ROIs to ensure that we only retain ROIs that show calcium-related activity. The ROIs that should be accepted are those that show at least one sizable peak or some significant slope (positive or negative), that hint at an activity. The ROIs that should be rejected are those that are too "flat".

The test that is applied at this step is the "mean absolute deviation" test (MAD). This test provides an idea of the variability of values in a dataset. The following process is used:

- Set the size of the window as configured using the value of `MEAN_ABS_DEVIATION_WINDOW` (the default configuration sets the value to 40);
- For each ROI data series in the data frame, compute the subtracted data series. A new data frame will be created that contains all the new ROI data series obtained by applying to each one of the original ROI data series the following:

- Start by computing roi[39] - roi[0] and store the computed value in a brand new data series;
- Shift downwards in the series by one index, that is, compute roi[40] - roi[1]. Append this new value to the new data series;
- Repeat the previous step until the last row of the ROI series is encountered.

At this point, an entirely new data frame has been crafted. For purely informational purposes, the program prints the plots of the newly computed sliding-subtraction High and Medium intensity ROIs (all the ROIs in the initial data frame since we still have not rejected any of them).

- Compute the mean intensity value for each ROI in the new data frame. We get a data series of ROI mean values;
- For each value in each ROI data series of the new data frame, compute the absolute value of (value - mean), that is compute |value - mean|. Store the computed value in place, so that at the end of the computation the new data frame has been refreshed with the new values;
- For each ROI, compute the mean of the new intensity values obtained at the previous step. This produces a data series with the mean absolute deviation values for each ROI;
- Multiply the values obtained at the previous step by the configured `MEAN_ABS_DEVIATION_FACTOR` value to obtain, for each ROI, the actual threshold value used at the next step;
- Iterate in each ROI's data series and check if any of the intensity values is greater than the value obtained at the previous step (that is, if the value is above the threshold). If so, increment a counter for the corresponding ROI;
- For each ROI, check how many intensity values matched the condition above (that is, check the counter value). If the counter value is greater or equal to the configured `CONDITION_MATCHING_MIN_POINT_COUNT` value, then the ROI is accepted, otherwise it is rejected.

At this point we have categorized all the ROIs from the initial data frame in *accepted* and *rejected* ROIs. The accepted ROIs did show significant calcium-related activity, while the rejected ones appeared to have too low an activity. It is now possible to perform new tasks, as described below.

##### Baseline subtraction

Baseline subtraction is based on the determination of the lowest point in the whole ROI. However, we have not retained a single lowest point of the trace but the mean value of a configurable number of points around that lowest point. The configuration parameter is `MIN_MEAN_INTENSITY_HALF_WINDOW`. If that value is set to 3, for example, then the ROI window that is used for the mean min intensity value would be computed with N = 2*3 + 1 = 7 ROI intensity values: three values above the actual lowest intensity point in the ROI series and three values below that point. The actual lowest intensity point is also accounted for in the mean value calculation. This can be represented like below, where the dots are ROI points left (above) or right (below) of the lowest intensity value (x) in the ROI series:

```
[ . . . x . . . ]
        |
        v
        lowest intensity value
```

There are two specific situations that must be evaluated:

- If the lowest intensity value is found in the very first values of the ROI series
- If the lowest intensity value is found in the very last values of the ROI series

In both cases above, the minimum mean intensity value is computed by using the N first ROI intensity values (first case) or the N last ROI intensity values (second case).

Once the minimum mean intensity value has been computed for each ROI, it gets subtracted from all the points in the ROI series.

##### Exporting the accepted ROIs as a comma-separated values file

Right after having performed the baseline subtraction, the accepted ROIs are exported to the `rois.csv` file.

##### Plotting of all the ROIs graphs

Now that we have performed the baseline subtraction on all the ROIs (accepted and rejected alike), we can plot the graphs for the user to assess the filtering quality. If the filtering appears imperfect, a number of configurable parameters can be tweaked to better it.

The graphs are automatically displayed, as shown in Fig. 4.1. The user may modify some aspects of the layout to ensure that all the labels and legends are properly displayed and visible. Once the desired modifications have been performed, the user closes the plot window and the graphs get automatically saved to three files of different formats : `rois-plots.`<png|svg|pdf>.

##### ROI mean intensity integrations

The integration of ROI intensity values is performed in two distinct modes:

- The whole ROI is integrated
- The ROI is integrated according to integration intervals (if any) configured using the `INTERVALS` configuration parameter in the `[INTEGRATION]` section of the `config.cfg` configuration file.

##### Whole ROI integrations

The integration is performed by computing the area under the curve at any given [n, n+1] point pair of the ROI. The mean of these two points intensities is compounded by the frame acquisition rate of the frame stack. That acquisition rate is configured using the `ACQ_RATE` parameter in the `config.cfg` configuration file.

For a given ROI series, the integration is performed as described below.

1. Define an `integration_value` variable set to 0
2. Compute `sum` = roi[0] + roi[1] and `mean` = `sum` / 2
3. Compute `local_integ` = `mean` * `ACQ_RATE`
4. Add `local_integ` to `integration_value`
5. Shift downwards the ROI series by 1 row and perform steps 2-4
6. Stop iterating in the ROI series when the back (bottom, or last) point is reached.

At the end of the above loop, `integration_value` contains the integration value of the whole ROI.

##### Interval-based ROI integrations

If the `INTERVALS` configuration parameter in the `[INTEGRATION]` section of the `config.cfg` configuration file is set to some valid value (see the configuration file shipped with the source code for an example), the integration described in the previous paragraph is performed by iterating only in the specified row ranges of the ROI.

The integrations are performed separately for three distinct cases:

- The High mean intensity ROIs
- The Medium mean intensity ROIs

- Both the High *and* Medium (Total) mean intensity ROIs

The integration values that are computed each time are the following:

- The integration value
- The mean integration value
- The standard deviation of the integration value

###### Statistical computations over the ROI integrations

Once all the integration values have been computed, some statistical computations are performed and reported in the self-explanatory `stats.csv` file.

###### ROI area calculations

The ROI area calculations are performed by first loading the contents of the `roi-high-area-xy.csv` and `roi-medium-area-xy.csv` files into a Pandas data frame.

One important configurable parameter in these calculations is the `FRAME_TOTAL_AREA` parameter that describes the full area of the frames in the frame stack. A number of statistical computations are performed from this parameter value and the areas described in the above two files. The results of these computations are output to the same self-explanatory `stats.csv` file as above.

##### 4.5.5 Inter-ROI Pearson correlations

Once all the processing work described above has been performed, the user might configure `post-prOccam` to perform a Pearson correlation between all the accepted ROIs.

The configuration key in the `config.cfg` file is `PERFORM_ROI_CORRELATIONS` in the `[CORRELATIONS]` section. If set to true, the inter-ROIs Pearson correlations are computed and the resulting matrix is displayed as shown in Fig. 4.2.

Another configuration key is `ROI_CORRELATION_THRESHOLD` that sets the threshold for ROIs to be considered correlated. Note that this is an absolute value, because correlations can be either positive (the correlation value bears a positive sign) or negative (the correlation value bears a negative sign). Thus, two ROIs are considered correlated if their correlation value is either less than -`ROI_CORRELATION_THRESHOLD` or greater than `ROI_CORRELATION_THRESHOLD`.

CHAPTER

FIVE

### POST-PROCCAM SUPERVISOR MANUAL

In the previous *post-prOccam* chapter, we described the Python software package that is used to post-process the output of the OCCAM Fiji/ImageJ2-based plugin.

The `pp-supervisor` Python script is designed to run automatically the post-prOccam software over all the data directories located in a given master directory.

The data file hierarchy needs to be organized in the following manner for the `pp-supervisor` program to run correctly:

```
master_data_dir/
                |
                |__ data_dir_1/
                |              |_ roi-high-mean-intensity.csv
                |              |_ roi-high-area.csv
                |              |_ roi-medium-mean-intensity.csv
                |              |_ roi-medium-area.csv
                |              |_ <optionally, config.cfg>
                |
                |
                |__ data_dir_2
                |              |_ roi-high-mean-intensity.csv
                |              |_ roi-high-area.csv
                |              |_ roi-medium-mean-intensity.csv
                |              |_ roi-medium-area.csv
                |              |_ <optionally, config.cfg>
                .
                .
                .
```

#### 5.1 Execution of the supervising script

##### 5.1.1 Automatic configuration of the `post-proccam` run

The `pp-supervisor` script is run from the command line. To get a help message, enter

`pp-supervisor -h` or `pp-supervisor --help`

There is a nice feature that allows one to create a reference `config.cfg` configuration file for the `post-proccam` run and to automatically distribute it to all the data directories. There are two ways to use the reference configuration file:

- Force copy of the reference configuration file to all the data directories:

  If the `pp-supervisor` program is run using the following command line:

```
</path/to/pp-supervisor> --force_config_copy <path/to/post-proccam> <path/to/
master_data_dir> <path/to/reference-config.cfg>
```

the `reference-config.cfg` will be copied to each data directory, irrespective of its previous existence in that directory.

- Only copy the reference configuration file to the data directories that have none:

  If the `pp-supervisor` program is run using the following command line:

  ```
  <path/to/pp-supervisor> <path/to/post-proccam> <path/to/master_data_dir> <path/to/
  reference-config.cfg>
  ```

  that is, without the `--force_config_copy` option, the `reference-config.cfg` will only be copied to data directories that do not contain one already.

The `pp-supervisor` program sequentially visits all the data directories contained in the master data directory and runs the following command:

```
<path/to/post-proccam> roi-high-mean-intensity.csv roi-high-area.csv
roi-medium-mean-intensity.csv roi-medium-area.csv config.cfg
```

##### 5.1.2 Output of the `pp-supervisor` script

Upon running the `post-proccam` program, any output from that program is printed to the console. If there is one error, the supervising script exits prematurely and outputs an error message.
