## Supplementary Figures for "Versatile and automated workflow for the analysis of oligodendroglial calcium signals in preclinical mouse models of myelin repair"

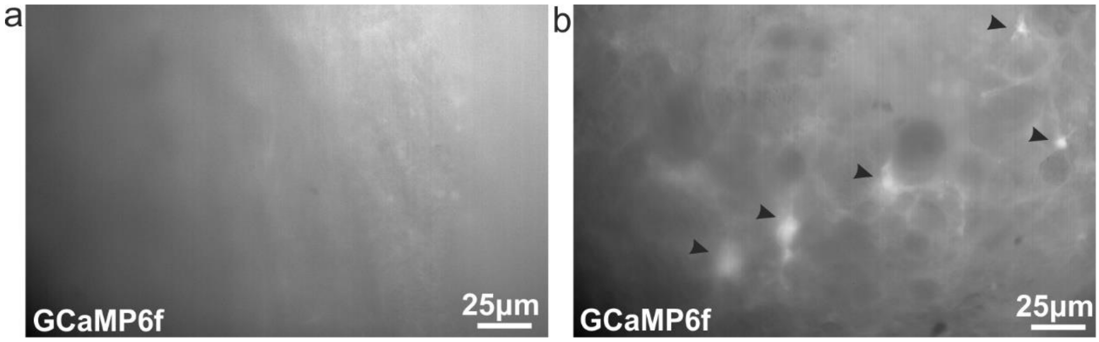


**Supplementary Figure 1 - GCaMP6f expression in non-lesioned versus lesioned corpus callosum. (a)** Representative image of GCaMP6f expression in the non-lesioned corpus callosum. Note that 4-hydroxy-tamoxifen injections were performed already in the second postnatal week and this image was taken at P55-65 (n=3). **(b)** Representative of GCaMP6f expression in the lesioned corpus callosum. Black arrows indicate GCaMP6f expressing ROIs. Here the hydroxy-tamoxifen injections were performed few days before the LPC injection as described in Figure 1.

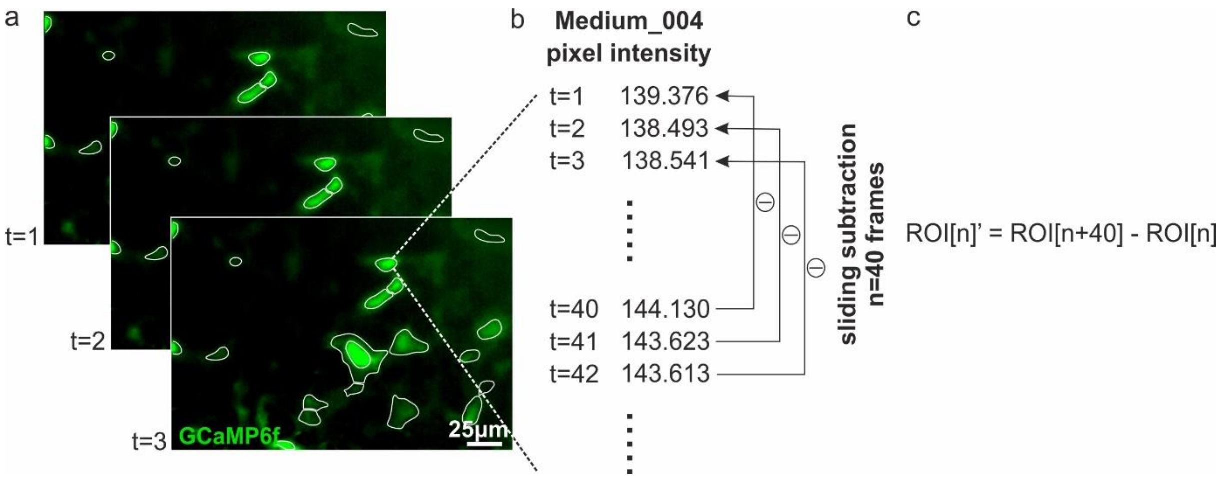


**Supplementary Figure 2 - ROI representation and sliding subtraction visualization. (a)** Schematic representation of a time resolved $Ca^{2+}$ imaging stack **(b)** Representation of raw data produced by Occam that is post-processed by post-prOccam including a time resolved list of mean pixel intensities for each ROI in the image stack. **(c)** A sliding subtraction method with formula ROI[n]' = ROI[n+40] - ROI[n] is performed on each ROI's mean pixel intensities at all timepoints and the resulting trace is used in a ROI rejection procedure that allows rejection of false positive ROIs. t: time, n: number of frames.

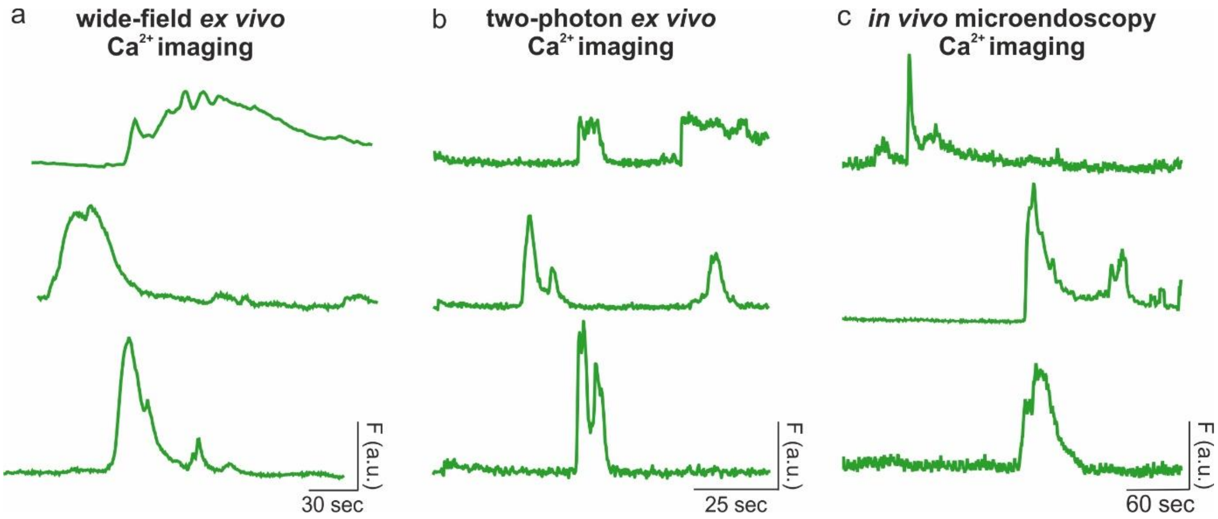


**Supplementary Figure 3 - Examples of complex oligodendroglial $Ca^{2+}$ signals.** Representative traces that include a complex $Ca^{2+}$ event with multiple peaks and lasting several seconds to several minutes. These complex $Ca^{2+}$ events are typical for oligodendroglia and can be observed in **(a)** wide-field ex vivo $Ca^{2+}$ imaging, **(b)** two-photon ex vivo $Ca^{2+}$ imaging and **(c)** in vivo microendoscopy $Ca^{2+}$ imaging.
