## Supplementary Files for "Versatile and automated workflow for the analysis of oligodendroglial calcium signals in preclinical mouse models of myelin repair"

### Supplementary File 1 - post-prOccam configuration file for wide field ex vivo imaging

```
[MAIN]

ACQ_RATE = 0.5716
FRAME_TOTAL_AREA = 432000

# The ROIs can be trimmed. To define trimming intervals uncomment the following
# lines and set appropriate values.  The "14 - LAST" format is a convenience
# format to trim the last 14 row of the ROIs. Change the 14 value to any
# appropriate value.

ROI_TRIM_INTERVALS =
  1-14
  #175-224
  14 - LAST

# The number of ROI points to be used to compute the
# minimum mean intensity of the ROI.
# The minimum mean intensivy is computed for the following window
# with MIN_MEAN_INTENSITY_HALF_WINDOW = 3:
#
#          [...|...]
#              v
#              min value
# 3 points are selected on the left the of minimum intensity value of the ROI
# and the same number of points are selected on the right of that minimum
# intensity value. The total window thus comprises 7 points.
MIN_MEAN_INTENSITY_HALF_WINDOW = 3

# The number of ROI points that are used to compute the ROI[z] - ROI[x]
# subtraction over all the ROI intensity values. If MEAN_ABS_DEVIATION_WINDOW =
# 40, the z-x window comprises 40 points.
MEAN_ABS_DEVIATION_WINDOW = 40

# Number by which the mean absolute deviation values are multiplied.
MEAN_ABS_DEVIATION_FACTOR = 1.5

# Number of points in each ROI that must have an intensity value greater than
# the threshold in order for the ROI to be accepted.
CONDITION_MATCHING_MIN_POINT_COUNT = 6

# Can be DEBUG or WARNING or INFO
DEBUG_OUTPUT_LEVEL = DEBUG

[INTEGRATION]
#
# A list of intervals (unit is the number of the frame
# in the image stack). To set intervals, uncomment the lines below and replace
```

```
# with your own values. Remove any unnecessary interval or add any required one.
# After the number interval, put a descriptive label inside double quotes.
# Please, do not comment out the "[INTEGRATION]" line nor the "INTERVALS ="
# line, as the program will fail if these lines are commented out or absent from
# this configuration file.
INTERVALS =
  1-362 "spontaneous"
  # 157-312 "after stim "
  #200-300 "wash"

[CORRELATIONS]

# Set the True if the inter-ROI correlations must be performed.
#PERFORM_ROI_CORRELATIONS = False
PERFORM_ROI_CORRELATIONS = True
# Threshold above which two ROIs are considered correlated.
# This is an absolute value. ROIs will be considered correlated if their
# correlation value is either less than -ROI_CORRELATION_THRESHOLD or greater
# than ROI_CORRELATION_THRESHOLD.
ROI_CORRELATION_THRESHOLD = 0.9
```

**Supplementary File 2 - post-prOccam configuration file for two-photon ex vivo imaging**

```
[MAIN]

ACQ_RATE = 0.2116
FRAME_TOTAL_AREA = 15732

# The ROIs can be trimmed. To define trimming intervals uncomment the following
# lines and set appropriate values.  The "14 - LAST" format is a convenience
# format to trim the last 14 row of the ROIs. Change the 14 value to any
# appropriate value.

ROI_TRIM_INTERVALS =
  1-2
  #175-224
  2 - LAST

# The number of ROI points to be used to compute the
# minimum mean intensity of the ROI.
# The minimum mean intensivy is computed for the following window
# with MIN_MEAN_INTENSITY_HALF_WINDOW = 3:
#
#          [...|...]
#              v
#              min value
# 3 points are selected on the left the of minimum intensity value of the ROI
# and the same number of points are selected on the right of that minimum
# intensity value. The total window thus comprises 7 points.
MIN_MEAN_INTENSITY_HALF_WINDOW = 9
# The number of ROI points that are used to compute the ROI[z] - ROI[x]
# subtraction over all the ROI intensity values. If MEAN_ABS_DEVIATION_WINDOW =
# 40, the z-x window comprises 40 points.
```

```
MEAN_ABS_DEVIATION_WINDOW = 15

# Number by which the mean absolute deviation values are multiplied.
MEAN_ABS_DEVIATION_FACTOR = 3

# Number of points in each ROI that must have an intensity value greater than
# the threshold in order for the ROI to be accepted.
CONDITION_MATCHING_MIN_POINT_COUNT = 4

# Can be DEBUG or WARNING or INFO
DEBUG_OUTPUT_LEVEL = DEBUG

[INTEGRATION]
#
# A list of intervals (unit is the number of the frame
# in the image stack). To set intervals, uncomment the lines below and replace
# with your own values. Remove any unnecessary interval or add any required one.
# After the number interval, put a descriptive label inside double quotes.
# Please, do not comment out the "[INTEGRATION]" line nor the "INTERVALS ="
# line, as the program will fail if these lines are commented out or absent from
# this configuration file.
INTERVALS =
  1-300 "CTRL"
  #150-250 "Carbachol "
  #200-300 "wash"

[CORRELATIONS]

# Set the True if the inter-ROI correlations must be performed.
#PERFORM_ROI_CORRELATIONS = False
PERFORM_ROI_CORRELATIONS = True
```

```
# Threshold above which two ROIs are considered correlated.
# This is an absolute value. ROIs will be considered correlated if their
# correlation value is either less than -ROI_CORRELATION_THRESHOLD or greater
# than ROI_CORRELATION_THRESHOLD.
ROI_CORRELATION_THRESHOLD = 0.9
```

**Supplementary File 3 - post-prOccam configuration file for in vivo microendoscopy imaging**

```
[MAIN]

ACQ_RATE = 0.2
FRAME_TOTAL_AREA = 80938

# The ROIs can be trimmed. To define trimming intervals uncomment the following
# lines and set appropriate values.  The "14 - LAST" format is a convenience
# format to trim the last 14 row of the ROIs. Change the 14 value to any
# appropriate value.

ROI_TRIM_INTERVALS =
  1-2
  #175-224
  2 - LAST

# The number of ROI points to be used to compute the
# minimum mean intensity of the ROI.
# The minimum mean intensivy is computed for the following window
# with MIN_MEAN_INTENSITY_HALF_WINDOW = 6:
#
#          [...|...]
#              v
#              min value
# 3 points are selected on the left the of minimum intensity value of the ROI
# and the same number of points are selected on the right of that minimum
# intensity value. The total window thus comprises 7 points.
MIN_MEAN_INTENSITY_HALF_WINDOW = 9

# The number of ROI points that are used to compute the ROI[z] - ROI[x]
# subtraction over all the ROI intensity values. If MEAN_ABS_DEVIATION_WINDOW =
# 40, the z-x window comprises 40 points.
MEAN_ABS_DEVIATION_WINDOW = 70

# Number by which the mean absolute deviation values are multiplied.
MEAN_ABS_DEVIATION_FACTOR = 5

# Number of points in each ROI that must have an intensity value greater than
# the threshold in order for the ROI to be accepted.
CONDITION_MATCHING_MIN_POINT_COUNT = 12

# Can be DEBUG or WARNING or INFO
DEBUG_OUTPUT_LEVEL = DEBUG

[INTEGRATION]
#
# A list of intervals (unit is the number of the frame
# in the image stack). To set intervals, uncomment the lines below and replace
# with your own values. Remove any unnecessary interval or add any required one.
# After the number interval, put a descriptive label inside double quotes.
# Please, do not comment out the "[INTEGRATION]" line nor the "INTERVALS ="
```

```
# line, as the program will fail if these lines are commented out or absent from
# this configuration file.
INTERVALS =
  1-2096 "spontaneous"
  # 157-312 "after stim "
  #200-300 "wash"

[CORRELATIONS]

# Set the True if the inter-ROI correlations must be performed.
#PERFORM_ROI_CORRELATIONS = False
PERFORM_ROI_CORRELATIONS = True
# Threshold above which two ROIs are considered correlated.
# This is an absolute value. ROIs will be considered correlated if their
# correlation value is either less than -ROI_CORRELATION_THRESHOLD or greater
# than ROI_CORRELATION_THRESHOLD.
ROI_CORRELATION_THRESHOLD = 0.9
```
